# Discovery of novel putative tumor suppressors from CRISPR screens reveals rewired lipid metabolism in AML cells

**DOI:** 10.1101/2020.10.08.332023

**Authors:** W. Frank Lenoir, Micaela Morgado, Peter C DeWeirdt, Megan McLaughlin, Audrey L Griffith, Annabel K Sangree, Marissa N Feeley, Nazanin Esmaeili Anvar, Eiru Kim, Lori L Bertolet, Medina Colic, Merve Dede, John G Doench, Traver Hart

**Affiliations:** The University of Texas MD Anderson Cancer Center UTHealth Graduate School of Biomedical Sciences; The University of Texas MD Anderson Cancer Center, Houston, TX; Department of Bioinformatics and Computational Biology, The University of Texas MD Anderson Cancer Center, Houston, TX, USA; Genetic Perturbation Platform, Broad Institute of MIT and Harvard, Cambridge, MA, USA; Department of Cancer Biology, The University of Texas MD Anderson Cancer Center, Houston, TX, USA

## Abstract

CRISPR knockout screens in hundreds of cancer cell lines have revealed a substantial number of context-specific essential genes that, when associated with a biomarker such as lineage or oncogenic mutation, offer candidate tumor-specific vulnerabilities for targeted therapies or novel drug development. Data-driven analysis of knockout fitness screens also yields many other functionally coherent modules that show emergent essentiality or, in rarer cases, the opposite phenotype of faster proliferation. We develop a systematic approach to classify these suppressors of proliferation, which are highly enriched for tumor suppressor genes, and define a network of 145 genes in 22 discrete modules. One surprising module contains several elements of the glycerolipid biosynthesis pathway and operates exclusively in a subset of AML lines, which we call Fatty Acid Synthesis/Tumor Suppressor (FASTS) cells. The proliferation suppressor activity of genes involved in the synthesis of saturated fatty acids, coupled with a more severe fitness phenotype for the desaturation pathway, suggests that these cells operate at the limit of their carrying capacity for saturated fatty acids, which we confirmed biochemically. Overexpression of genes in this module is associated with a survival advantage in an age-matched cohort of AML patients, suggesting the gene cluster driving an *in vitro* phenotype may be associated with a novel, clinically relevant subtype.

## Introduction

Gene knockouts are a fundamental tool for geneticists, and the discovery of CRISPR-based genome editing^1^ and its adaptation to gene knockout screens has revolutionized mammalian functional genomics and cancer targeting^2–8^. Hundreds of CRISPR/Cas9 knockout screens in cancer cell lines have revealed background-specific genetic vulnerabilities^9–13^, providing guidance for tumor-specific therapies and the development of novel targeted agents. Although lineage and mutation state are powerful predictors of context-dependent gene essentiality, variation in cell growth medium and environment can also drive differences in cell state, particularly among metabolic genes^14,15^, and targeted screening can reveal the genetic determinants of metabolic pathway buffering^16,17^.

The presence and composition of metabolic and other functional modules in the cell can also be inferred by integrative analysis of large numbers of screens. Correlated gene knockout fitness profiles, measured across hundreds of screens, have been used to infer gene function and the modular architecture of the human cell^18–21^. Data-driven analysis of correlation networks reveals clusters of functionally related genes whose emergent essentiality in specific cell backgrounds is often unexplained by the underlying lineage or mutational landscape^21^. Interestingly, in a recent study of paralogs whose functional buffering renders them systematically invisible to monogenic CRISPR knockout screens^22,23^, it was shown that the majority of context-dependent essential genes are constitutively expressed in cell lines^23^. Collectively these observations suggest that there is much unexplained variation in the genetic architecture, and emergent vulnerability, of tumor cells.

Building human functional interaction networks from correlated gene knockout fitness profiles in cancer cells is analogous to generating functional interaction networks from correlated genetic interaction profiles in *S. cerevisiae*^24–27^. The fundamental difference between the two approaches is that, in yeast, a massive screening of pairwise gene knockouts in a single yeast strain was conducted in order to measure genetic interaction - a dual knockout phenotype more or less severe than that expected by the combination of the two genes independently. In coessentiality networks, CRISPR-mediated single gene knockouts are conducted across a panel of cell lines that sample the diversity of cancer genotypes and lineages. Digenic perturbations in human cells, a more faithful replication of the yeast approach, are possible with Cas9 and its variants, but library construction, sequencing, and positional biases can be problematic^16,28–34^. Recently, we showed that an engineered variant of the Cas12a endonuclease, enCas12a^35^, could efficiently perform multiplex gene knockouts^34^, and we demonstrated its effectiveness in assaying synthetic lethality between targeted paralogs^23^. These developments in principle enable researchers to measure how biological networks vary across backgrounds, a powerful approach for deciphering complex biology^24,36,37^.

CRISPR perturbations in human cells can result in loss of function alleles that increase as well as decrease *in vitro* proliferation rates; faster proliferation is an extreme rarity in yeast knockouts. These fast-growers can complicate predictions of genetic interaction^29^ and confound pooled chemoresistance screens^38^. However, there is no broadly accepted method of identifying these genes from CRISPR screens. Here we describe the development of a method to systematically classify genes whose knockout provides a proliferation advantage *in vitro*. We observe that genes which confer proliferation advantage are typically tumor suppressor genes, and that they show the same modularity and functional coherence as context-dependent essential genes. Moreover, we discover a novel module that includes several components of the glycerolipid biosynthesis pathway that slows cell proliferation in a subset of acute myeloid leukemia (AML) cell lines. We show a rewired genetic interaction network using enCas12a multiplex screening, and find strong genetic interactions corroborated by clinical survival data. A putative tumor-suppressive role for glycerolipid biosynthesis is surprising and disconcerting, since this process is thought to be required to generate biomass for tumor cell growth, and inhibitors targeting this pathway are currently in clinical trials^39,40^.

## Results

### Identifying Proliferation Suppressor Signatures

We previously observed genes whose knockout leads to overrepresentation in pooled library knockout screens. These genes, which we term proliferation suppressor genes (PSG), exhibit positive selection in fitness screens, a phenotype opposite that of essential genes. As expected, many PSG are known tumor suppressor genes; for example, *TP53* and related pathway genes *CDKN1A*, *CHEK2*, and *TP53BP1* show positive selection in select cell lines (**Figure 1a**). Detection of these genes as outliers is robust to the choice of CRISPR analytical method, as we tested BAGEL2^41,42^, CERES^10^, JACKS^43^, and mean log fold change (LFC) of gRNA targeting each gene (**Supplementary Figure 1a-d**). Unlike core-essential genes, PSG are highly context-specific: *TP53* knockout shows positive LFC only in cell lines with wild-type *TP53* (**Figure 1b**), and *PTEN* knockout shows the PS phenotype only in *PTEN^wt^* backgrounds (**Figure 1c**). These observations are consistent with the knockout phenotypes of known tumor suppressor genes (TSG) in cell lines: in wildtype cells, TSG knockout increases the proliferation rate in cell culture, but when cell lines are derived from tumors where the TSG is already lost or non-functional, gene knockout has no effect. TSG are therefore context-specific PSG, but it is not necessarily the case that genes with a proliferation suppressor phenotype *in vitro* act as TSG *in vivo;* proliferation suppressors are at best putative tumor suppressors in the absence of confirmatory data from tumor profiling.

**Figure 1.**
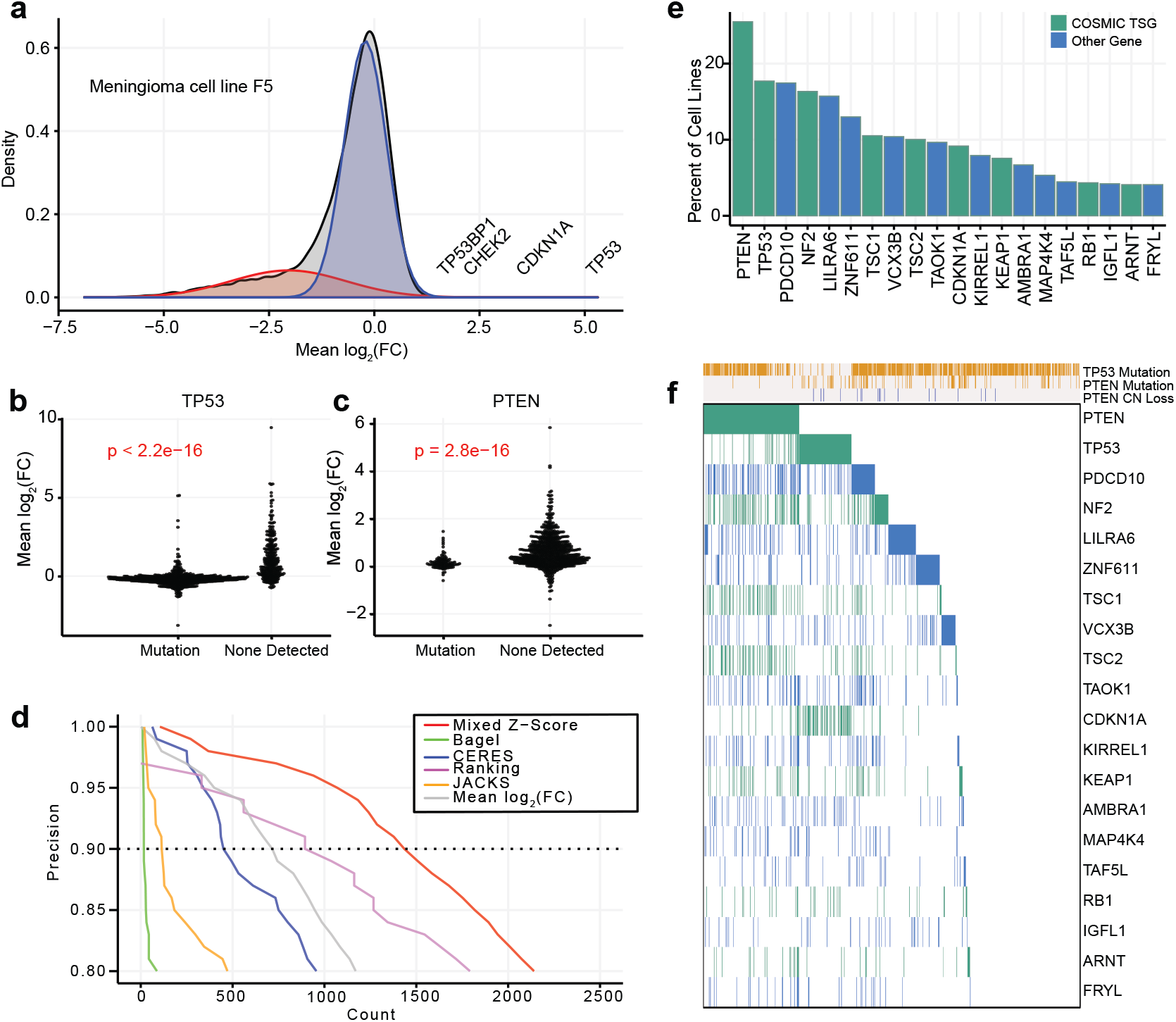
Discovery of Proliferation Suppressor genes. (a) Fold-change distribution of a typical CRISPR knockout screen has a long left tail of essential genes, and a small number of genes whose knockout increases fitness (proliferation suppressor genes, “PSG”). A two-component Gaussian mixture model (red, blue) models this distribution. (b) and (c) Fold change of common tumor suppressors across 808 cell lines (P-values, Wilcoxon rank-sum tests). (d) Precision vs. recall of mixed Z-score and other CRISPR analysis methods. Dashed line, 90% precision (10% FDR). (e) Fraction of cell lines in which known tumor suppressors (green) or other genes (blue, not defined as TSG by COSMIC) are classified as PS genes at 10% FDR. (f) Presence of each known TSG across 808 cell lines, vs. cell genetic background. Gold, mutation present; gray, absent. Green or blue, following color scheme in (e), gene is classified as a proliferation suppressor.

Though detection of PSG is possible using existing informatics pipelines, several factors complicate a robust detection of these genes. There is no accepted threshold for any algorithm we considered to detect PSG, since all were optimized to classify essential genes. A related second issue is that cell line screens show a wide range of variance in LFC distributions, making robust outlier detection challenging (**Supplementary Figure 1e-f**). Third, the signatures are strongly background-dependent, as demonstrated by PTEN and TP53. Finally, there is no consistent expectation for whether or how many putative tumor suppressor genes are present in a given cell line.

To address this gap, we developed a method to account for variability in fold-change distributions between screens. Our approach uses a Gaussian mixture model (K=2) to estimate each screen’s distribution of gene-level LFC scores (**Figure 1a**). Mixed distribution models have previously been used to identify distinctions between populations of essential and nonessential fitness genes in CRISPR screens^44^. For the *K* = 2 mixture model, the more negative distribution (**Figure 1a**, red) is generally essential genes, while the higher, narrower peak around zero (**Figure 1a**, blue), models the large population of knockouts with no fitness phenotype. We used this second model to calculate a Z-score (hereafter referred to as the ‘mixed Z-score’) for all gene-level mean fold changes in each cell line. This approach normalizes variance (**Supplementary Figure 1e-f**) across LFC distributions in different cell lines, with negative Z-scores indicating essential genes and positive Z-scores representing PSG phenotypes.

To evaluate the effectiveness of this mixed Z-score approach, we used COSMIC^45,46^ tumor suppressor genes as a true positive reference set, and we combined COSMIC-defined oncogenes (removing dual-annotated tumor suppressors) with our previously-specified set of nonessential genes as a true negative reference set^7,47^. Since there is no expectation for the presence of a consistent set of PSG across cell lines, we analyzed each of the 808 cell lines from the Avana 2020Q4 data release independently^10,48,49^ calculating gene-level scores on each cell line individually and then combining all scores into a master list of 808 x 18k = 14.6 million gene-cell line observations (**Supplementary Table 1**). Moreover, since there is also no expectation that all COSMIC TSG would be detected cumulatively across all cell lines, we judged that traditional recall metrics (e.g. percentage of the reference set recovered) were inappropriate. We therefore defined recall as the total number of TSG-cell line observations. Using this evaluation scheme, the mixed Z-score approach outperforms comparable methods by a substantial margin, classifying more than 722 PS-cell line instances at a 10% false discovery rate (FDR) (**Figure 1d**). This is roughly 50% more putative PSG than the closest alternative, a nonparametric rank-based approach, at the same FDR. BAGEL^41,42^, a supervised classifier of essential genes, performed worst at TSG, and the raw mean LFC approach also fared poorly, highlighting the need for variance normalization across experiments. We applied this 10% FDR threshold for all subsequent analyses.

Common tumor suppressor genes PTEN and TP53 were observed in ∼15% of cell lines (**Figure 1e**), with other well-known TSG appearing less frequently. Among 309 COSMIC TSGs for which we have fitness profiles (representing 1.7% of all 18k genes), we find that 116 (37.5%) of these genes occur as proliferation suppressors at least once (**Supplementary Table 2**) and make up 24.4% of total proliferation suppressor calls (**Supplementary Figure 2a-b**), a 14-fold enrichment. All of the known TSG hits come from just 504 of the 808 cell lines (62.4%) in which proliferation suppressor hit calls were identified (**Figure 1f**), yet we did not observe a bias toward particular tissues: in every lineage, most cell lines carried at least one PSG (**Supplementary Figure 1g**).

To further validate our approach, we compared the set of TSGs in our PSG hits to other molecular profiling data. When identified as a proliferation suppressor, 53% of the 116 TSGs demonstrate higher mean mRNA expression relative to backgrounds where the same TSG is not a proliferation suppressor (**Supplementary Table 2**). Similarly, 96.6% of the 116 TSGs, when classified as a proliferation suppressor, demonstrate lower frequency of nonsilent mutations compared to backgrounds where the TSG is not a hit (**Supplementary Table 2**). These observations were not restricted to COSMIC TSGs however, as this was the case for all PSG hit calls of genes against non-PSG hit calls (**Supplementary Figure 2c** and **2d**). Copy number comparisons did not suggest major distinctions between PSG vs non-PSG calls (**Supplementary Figure 2e)**, however there did appear to be more variation in PSG observations, possible stemming from smaller grouped sample sizes. Together, these observations confirm the reliability of our method to detect genes whose knockout results in faster cell proliferation, and that, analogous to essential genes, these genes must be expressed and must not harbor a loss-of-function mutation in order to elicit this phenotype.

We attempted to corroborate our findings using a second CRISPR dataset of 342 cell line screens from Behan *et al.*^13^, including >150 screens in the same cell lines as in the Avana data. However, these screens were conducted over a shorter timeframe than the Avana screens (14 vs. 21 days), giving less time for both positive and negative selection signals to appear (see Methods for a detailed discussion). As a result, when we compared cell lines screened by both groups, the Avana data yielded many more TSG hits (**Supplementary Figure 3a**). While most of these do not meet our threshold for PSG in the Sanger data, hits at our 10% FDR threshold across all Avana screens are strongly biased toward positive mixed Z-scores in the Sanger screens (**Supplementary Figure 3b**), consistent with a weaker signal of positive selection as a result of the shorter assays rather than a lack of robustness in the screens^49^.

### Discovering Pathways Modulating Cell Growth with a Proliferation Suppressor Co-Occurrence Network

Although known TSG act as PSG in only a subset of cell lines, we observed patterns of co-occurrence among functionally related genes. *PTEN* co-occurs with mTOR regulators *NF2*^50^ (P < 3×10^−11^, Fisher’s exact test) and the *TSC1*/*TSC2* complex (P-values both < 7×10^−13^)^51^, as well as Programmed Cell Death 10 (*PDCD10*)^52^, a proposed tumor suppressor^7,53^ (**Figure 2a**). The p53 regulatory cluster (*TP53*, *CDKN1A*, *CHECK2*, *TP53BP1*) also exhibited a strong co-occurrence pattern that was independent of the mTOR regulatory cluster (**Figure 2a**). mTOR^54^ and cell cycle checkpoint genes^55,56^ have been heavily linked to cancer development, given their roles in controlling cell growth and proliferation, and thus have been the focus of studies characterizing patient genomic profiles to identify common pathway alterations^57,58^.

**Figure 2.**
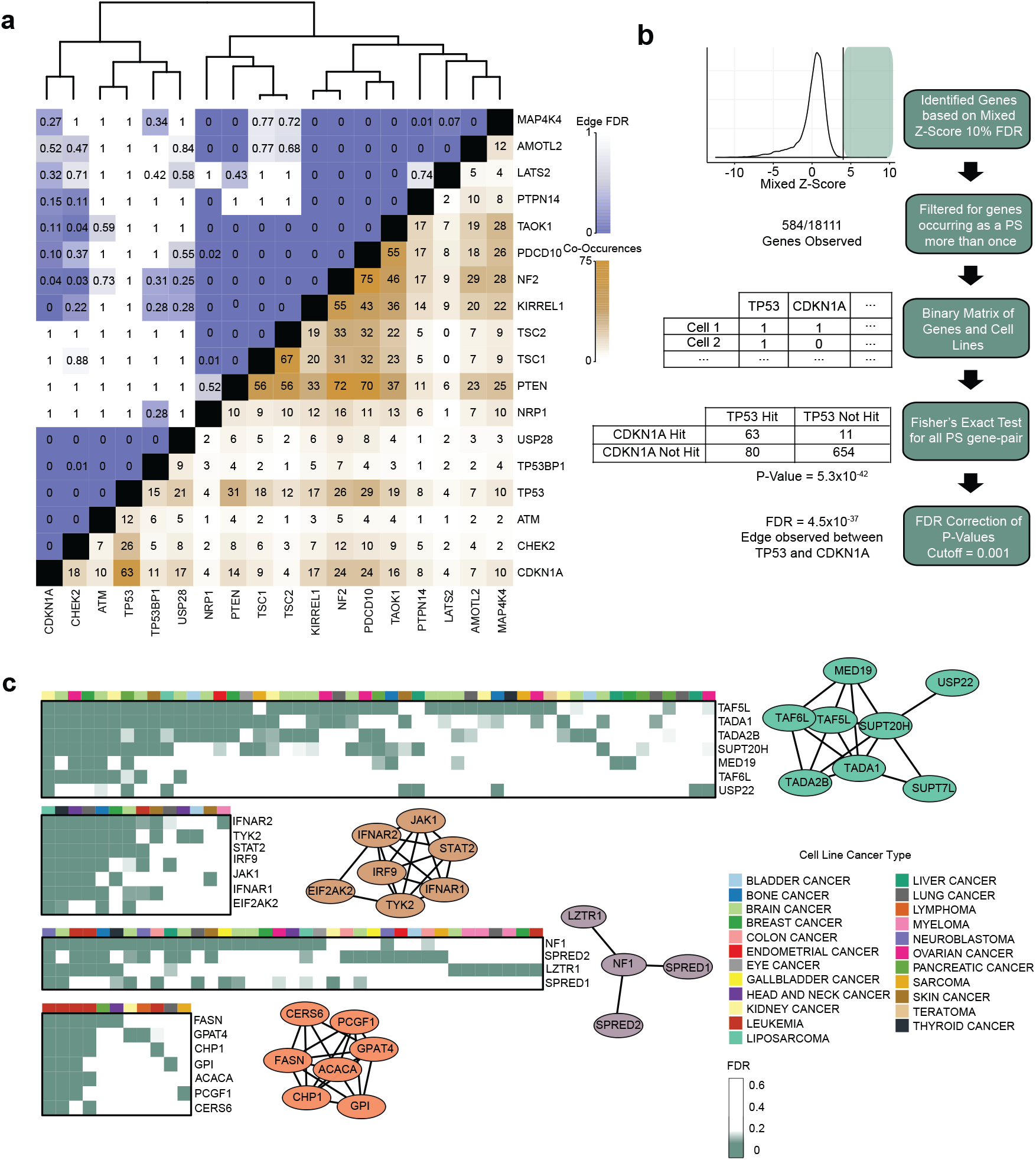
Co-occurrence of PSG. (a) Co-occurrence/mutual exclusivity of common TSG as PSG in CRISPR screens. Brown, number of cell lines in which two genes co-occur as PSG at 10% FDR. Blue, FDR of co-occurrence. Hierarchical clustering delineates functional modules. (b) Pipeline for building the co-PS network. (c) Examples from the Co-PS network. Nodes are connected by edges at FDR < 0.1%. Heatmaps indicate presence of PSG vs. cell lineage.

The modularity of mTOR regulators and TP53 regulators demonstrates pathway-level proliferation suppressor activity. This reflects the concept of genes with correlated fitness profiles indicating the genes operate in the same biochemical pathway or biological process^19,21,59,60^. However, the sparseness of PSG, coupled with their smaller effect sizes, renders correlation networks relatively poor at identifying modules of genes with proliferation suppressor activity. In order to identify such modules, we developed a PSG network (**Supplementary Table 3**) based on statistical overrepresentation of co-occurring PSG (**Figure 2b**); see Methods for details. This approach yields a network of 145 genes containing 462 edges in disconnected clusters; only 8 clusters have 3 or more genes (**Figure 2c** and **Supplementary Figure 4c**). Of these 462 edges, 74 (16.0%, empirical P<10^−4^) are present in the HumanNet^61^ functional interaction network (**Supplementary Figure 4a-b**),∼8 fold more than expected from random sampling, indicating high functional coherence between connected genes. The network recovers the PTEN and TP53 modules as well as the Hippo pathway, the aryl hydrocarbon receptor complex (AHR/ARNT), the mTOR-repressing GATOR1 complex, the STAGA chromatin remodeling complex, JAK-STAT signaling, and the gamma-secretase complex (**Figure 2c, and Supplementary 4c**), all of which have been associated with tumor suppressor activity. The functional coherence and biological relevance of the PSG co-occurrence network further validates the approach taken and establishes this dataset as a resource for exploring putative tumor suppressor activity in cell lines and tumors.

### Variation in Fatty Acid Metabolism in AML Cells

In addition to the known tumor suppressors, we observed a large module containing elements of several fatty acid (FA) and lipid biosynthesis pathways (**Figure 2c**). Interestingly, while there does not appear to be a strong tissue specificity signature for most clusters (**Figure 2c**), the fatty acid metabolism cluster shows a strong enrichment for AML cell lines (P = 1.5×10^−4^). AML, like most cancers, typically relies on increased glucose consumption for energy and diversion of glycolytic intermediates for the generation of biomass required for cell proliferation. Membrane biomass is generated by phospholipid biosynthesis that uses fatty acids as building blocks, with FA pools replenished by some combination of triglyceride catabolism, transporter-mediated uptake, and *de novo* synthesis via the *ACLY/ACACA/FASN* palmitate production pathway using citrate precursor diverted from the TCA cycle. Indeed, the role of lipid metabolism in AML progression is indicated by changes in serum lipid content^62^, in particular for long-chain saturated fatty acids that are the terminal product of the FAS pipeline. Inhibition of FA synthesis is therefore an appealing chemotherapeutic intervention^63,64^ and FASN inhibitors are currently undergoing clinical trials for treatment of solid tumors and metabolic diseases^40^. The observation that knocking out FAS pathway genes results in *faster* proliferation in some AML cells, and their signature as putative tumor suppressor genes, is therefore very unexpected.

To learn whether additional elements of lipid metabolism were associated with the FAS cluster, we examined the differential correlation of mixed Z-scores in AML cells. We and others have shown that genes with correlated gene knockout fitness profiles in CRISPR screens are likely to be involved in the same biological pathway or process (“co-functional”)^18–21^, analogous to correlated genetic interaction profiles in yeast^25,26,65^. Strikingly, all gene pairs within the fully connected clique in the FAS cluster (containing genes *FASN*, *ACACA*, *GPAT4*, *CHP1*, *GPI CERS6, PCGF1,* **Figure 2c**) had a median Pearson correlation coefficient (PCC) of 0.76 in the 23 AML cell lines (range 0.63-0.95, **Figure 3a**, red), compared to median correlation of 0.05 in the remaining 785 cell lines (range -0.11-0.62, with the highest correlation between *FASN* and *ACACA*, adjacent enzymes in the linear palmitate synthesis pathway; **Figure 3a,** gray). These high differential Pearson correlation coefficients (dPCC) suggest that variation in lipid metabolism is pronounced in AML cells^66^.

**Figure 3.**
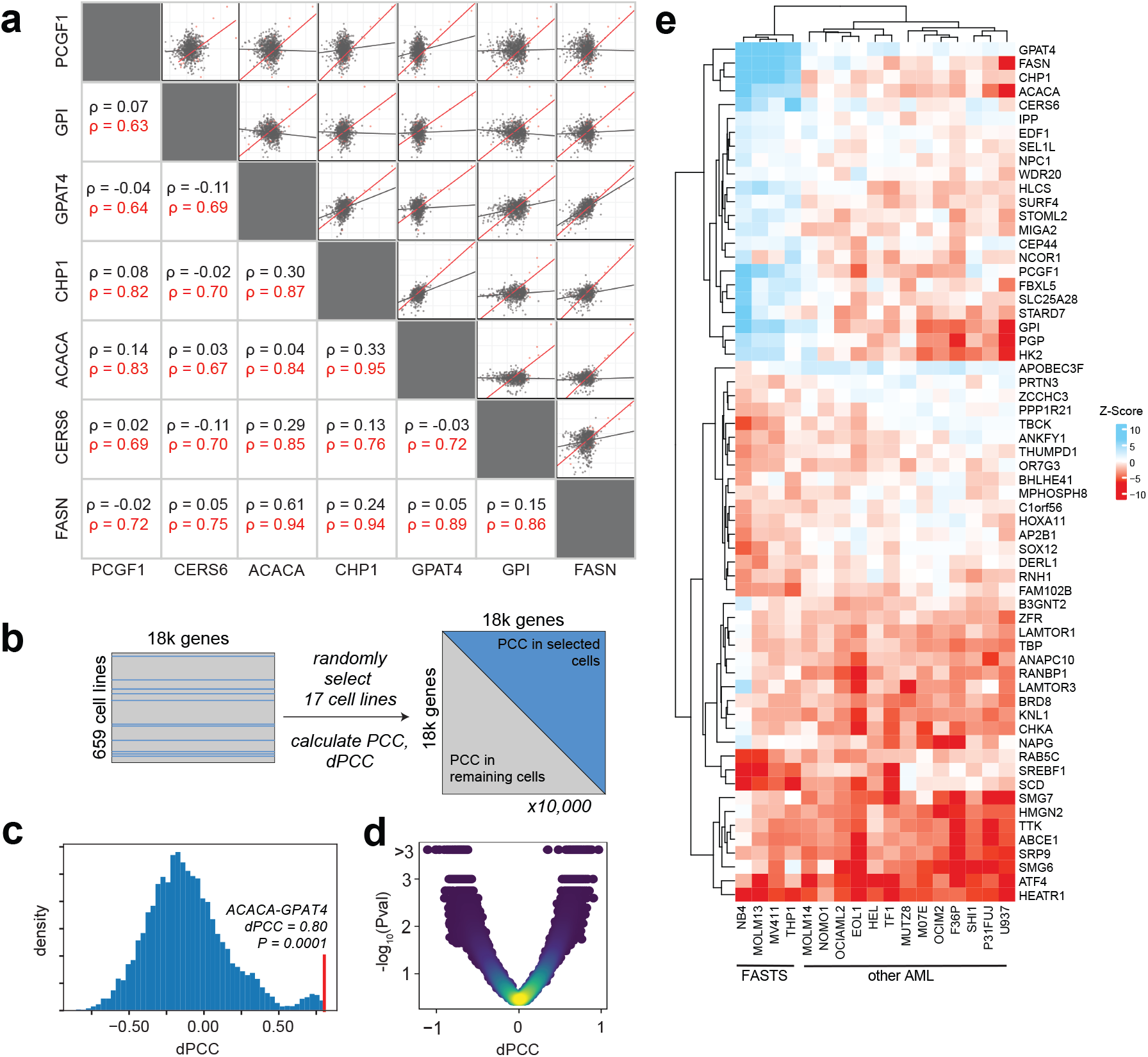
Differential network analysis of fatty acid synthesis module. (a) Among genes in the FAS module, Pearson correlation coefficients of shuffled Z score profiles are substantially higher in AML cells (red) than in other cells (gray). (b) Significance testing of differential PCC (dPCC) involves quality filtering of Avana data (n=659 cell lines, including 17 AML cell lines), building a null distribution by randomly selecting 17 cell lines, and calculating PCC between all gene pairs in the selected cells and the remaining cells. (c) After 1,000 repeats, a null distribution is generated for each pair, and a two-sided P-value is calculated for the observed AML-vs-other dPCC. (d) Volcano plot of dPCC vs. P-value for all genes in the Co-PS cluster. (e) Heatmap of mixed Z score for 17 AML cell lines in selected genes with high |mixed Z| and high |dPCC|. Clustering of cell lines indicates the putative Fatty Acid Synthesis/Tumor Suppressor (FASTS) subtype.

We sought to explore whether this difference in correlation identified other genes that might give insight into metabolic rewiring in AML. We first removed noisy data by filtering for high-quality screens (Cohen’s D > 2.5, recall > 60%^42^), leaving 659 cell lines, including 17 AML cell lines. Calculating a global difference between PCC of all gene pairs in all 17 AML and in the remaining 642 cell lines yielded many gene pairs whose dPCC appeared indistinguishable from random sampling (**Supplementary Figure 5a-b**). To filter these, we calculated empirical P-values for each gene pair. We randomly selected 17 cell lines from the pool of all screens, calculated PCC for all gene pairs in the selected and remaining lines, and calculated dPCC from these PCC values (**Figure 3b**). We repeated this process 1,000 times to generate a null distribution of dPCC values for each gene pair, against which a P-value could be computed (**Figure 3c-d**).

Expanding the set to a filtered list of genes whose correlation with a gene in the FAS clique showed significant change in AML cells (P<0.001; see Methods) yielded a total of 106 genes, including the 7 genes in the clique (**Figure 3e**) plus Holocarboxylase Synthetase (*HLCS*), which biotinylates and activates acetyl-CoA-carboxylase, the protein product of *ACACA*, as well as glycolysis pathway genes *PGP* and *HK2*. Interestingly, about half of the genes showed significantly increased anticorrelation with the FAS cluster, indicating genes preferentially essential where the FAS genes act as proliferation suppressors (**Figure 3e**). These genes include fatty acid desaturase (*SCD*), which operates directly downstream from *FASN/ACACA* to generate monounsaturated fatty acid species, and Sterol Regulatory Element Binding Transcription Factor 1 (*SREBF1*), the master regulatory factor for lipid homeostasis in cells.

Clustering the AML cells lines according to these high-dPCC genes reveals two distinct subsets of cells. The FAS cluster and its correlates show strong proliferation suppressor phenotype in four cell lines, NB4, MV411, MOLM13, and THP1. The remaining thirteen AML cell lines show negligible to weakly essential phenotypes when these genes are knocked out. The anticorrelated genes, including *SCD* and *SREBF1*, show heightened essentiality in these same cell lines. Together these observed shifts in gene knockout fitness indicates that this subset of AML cells has a substantial metabolic rewiring. Because these cells share a genetic signature among fatty acid synthesis pathway genes that is consistent with tumor suppressors, we call these cell lines <u>F</u>atty <u>A</u>cid <u>S</u>ynthesis/<u>T</u>umor <u>S</u>uppressor (FASTS) cells (**Figure 3e**).

### Cas12a-mediated Genetic Interaction Screens Confirm Rewired Lipid Metabolism

We sought to confirm whether gene knockout confers improved cell fitness, and to gather some insight into why some AML cells show the FASTS phenotype and others do not. Genetic interactions have provided a powerful platform for understanding cellular rewiring in model organisms, and we sought to apply this approach to deciphering the FASTS phenotype. We designed a CRISPR screen that measures the genetic interactions between eight selected “query genes” and ∼100 other genes (“array genes”). The query genes include *FASN* and *ACACA*, from the cluster of proliferation-suppressor genes, as well as lipid homeostasis transcription factor *SREBF1*, anticorrelated with the FAS cluster in the differential network analysis, and uncharacterized gene *c12orf49*, previously implicated in lipid metabolism by coessentiality^21^ and a recent genetic interaction study^60^. Additional query genes include control tumor suppressor genes *TP53* and *PTEN* and control context-dependent essential genes *GPX4* and *PSTK (***Figure 4a**). The array genes include two to three genes each from several metabolic pathways, including various branches of lipid biosynthesis, glycolysis and glutaminolysis, oxphos, peroxisomal and mitochondrial fatty acid oxidation. We include the query genes in the array gene set (**Figure 4a**) to test for screen artifacts and further add control essential and nonessential genes to measure overall screen efficacy (**Supplementary Table 4-5**).

**Figure 4.**
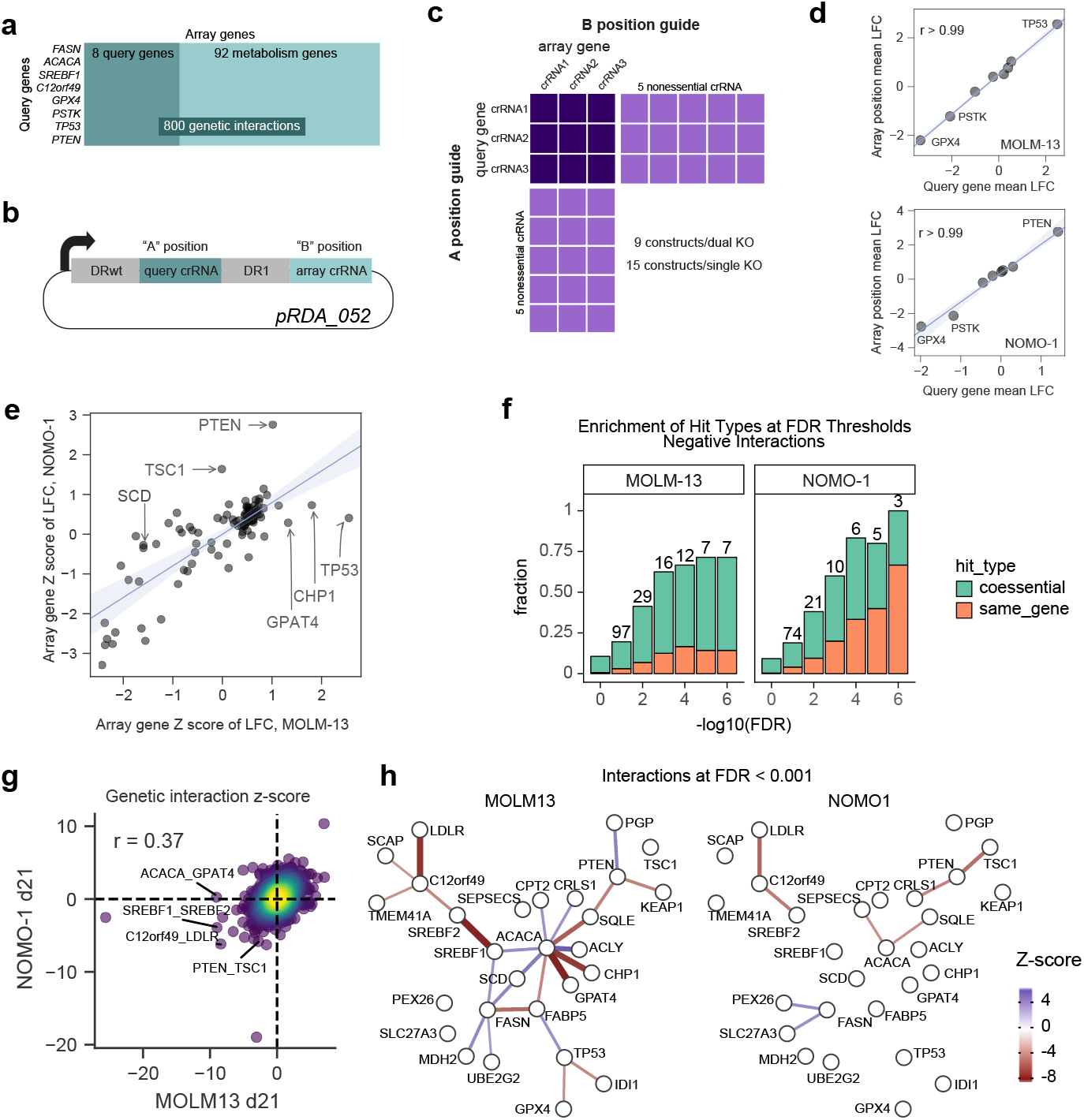
Genetic interactions reveal a rewired lipid biosynthesis pathway in FASTS cells. (a) Genetic interaction screen targets 8 query genes, selected from FASTS cluster and dPCC analysis, and 100 array genes sampling lipid metabolism pathways, for a total of 800 pairwise knockouts. (b) Library design uses a dual-guide enCa12a expression vector which targets the query gene in the “A” position and array gene in the “B” position. (c) Overall library design includes three crRNA/gene plus control crRNA targeting nonessential genes. Single-knockout constructs (target gene paired with nonessential controls) allow accurate measurement of single knockout fitness. (d) Considering single knockout fitness of query genes in the “A” and “B” position of the crRNA expression vector shows no position effects in the two cell lines screened (MOLM13, NOMO1). LFC, log fold change. (e) Single knockout fitness (Z-score of mean LFC) is highly consistent between MOLM13 and NOMO1, but reveals background-specific PS genes. (f) Enrichment among GI for coessential and self-interacting genes. Self-interactions among genes that show single knockout fitness phenotypes are expected, reflecting quality of GI observations. (g) Global comparison of MOLM13, NOMO1 genetic interaction Z scores. (h) Network view of interactions in each background shows rewiring in MOLM13 FASTS cells.

We used the enCas12a CRISPR endonuclease system to carry out multiplex gene knockouts^35^. We used a dual-guide enCas12a design, as described in DeWeirdt *et al.*^34^, that allows for construction of specific guide pairs through pooled oligonucleotide synthesis (**Figure 4b**). The library robustly measures single knockout fitness by pairing three Cas12a crRNA per target gene each with five crRNA targeting nonessential genes^7,47^ (n=15 constructs for single knockout fitness), and efficiently assays double knockout fitness by measuring all guides targeting query-array gene pairs (n=9) (**Figure 4c & Supplementary Table 5**). Using this efficient design and the endogenous multiplexing capability of enCas12a, we were able to synthesize a library targeting 800 gene pairs with a single 12k oligonucleotide array.

We screened one AML cell line from the FASTS subset, MOLM13, and a second one with no FAS phenotype, NOMO1, collecting samples at 14 and 21 days after transduction with a five-day puromycin selection (**Supplementary Table 6-7**). Importantly, by comparing the mean log fold change of query gene knockouts in the “A” position *vs.* the same genes in the “B” position of the dual knockout vector, we find no positional bias in the multiplex knockout constructs (**Figure 4d**), consistent with our previous findings^23,34^. Single knockout fitness measurements effectively segregated known essential genes from nonessentials, confirming the efficacy of the primary screens (**Supplementary Figure 6**). Context-dependent fitness profiles are consistent with the cell genotypes, with *PTEN* and *TSC1* showing positive selection in *PTEN^wt^* NOMO1 cells and *TP53* being a strong PS gene in *P53^wt^* MOLM13 cells. Strikingly, *CHP1* and *GPAT4* are the next two top hits in MOLM13, confirming their proliferation suppressor phenotype (**Figure 4e**), while neither shows a phenotype in NOMO1. Together these observations validate the enCas12a-mediated multiplex perturbation platform, confirm the ability of CRISPR knockout screens to detect proliferation suppressors, and corroborate the background-specific fitness enhancing effects of genes from the FAS cluster.

To measure genetic interactions, we fit a linear regression for each guide between the combination LFCs and the single guide LFCs, Z-scoring the residuals from this line, and combining across all guides targeting the same gene pair **(Supplementary Figure 6 & Supplementary Table 7**). Here, positive genetic interaction Z-scores reflect greater fitness than expected and negative Z-scores represent lower than expected based on the single gene knockouts independently, similar to the methodology applied in a recent survey of genetic interactions in cancer cells using multiplex CRISPR perturbation^33^ (see Methods). Gene self-interactions (when the same gene is in the A and B position, **Figure 4d**) should therefore be negative for proliferation suppressors and positive for essentials (**Figure 4f-g**, **Supplementary Figure 6**). Overall, genetic interaction Z-scores in the two cell lines showed moderate correlation (**Figure 4g**) and previously reported synthetic interactions between *C12orf49* and low-density lipoprotein receptor *LDLR*^17^ and between *SREBF1* and its paralog *SREBF2*^17^ are identified in both cell lines (**Supplementary Figure 6f-g**).

In contrast with the interactions found in both cell lines, background-specific genetic interactions reflect the genotypic and phenotypic differences between the cells. The negative interaction between tumor suppressor *PTEN* and mTOR repressor *TSC1* in *PTEN^wt^* NOMO1 cells is consistent with their epistatic roles in the mTOR regulatory pathway. Both genes show positive knockout fitness in NOMO1 (**Figure 4e**) but their dual knockout does not provide an additive growth effect, resulting in a suppressor interaction with a negative Z-score (**Figure 4g-h**). Similarly, suppressor genetic interactions between *ACACA* and downstream proliferation suppressor genes *CHP1* and *GPAT4* are pronounced in MOLM13 cells, consistent with epistatic relationships in a linear biochemical pathway (**Figure 4h**). These interactions are not replicated with query gene *FASN*, but both *FASN* and *ACACA* show negative interactions with fatty acid transport gene *FABP5* and positive interactions with *SREBF1* and *SCD,* the primary desaturase of long-chain saturated fatty acids. All of these interactions are absent in NOMO1, demonstrating the rewiring of the lipid biosynthesis genetic interaction network between these two cell types (**Figure 4h**).

### FASTS Signature Predicts Sensitivity to Saturated Fatty Acids

The significant differences in the single- and double-knockout fitness signatures between the two cell lines suggests a major rewiring of lipid metabolism in these cells. *CHP1* and *GPAT4* are reciprocal top correlates in the Avana coessentiality network (r = 0.43, P = 2.5×10^−34^), strongly predicting gene co-functionality^21^. Two recent studies characterized the role of lysophosphatidic acid acyltransferase *GPAT4* in adding saturated acyl moieties to glycerol 3-phosphate, generating lysophosphatidic acid (LPA) and phosphatidic acid (PA), the precursors for cellular phospholipids and triglycerides, and further discovered *CHP1* as a key regulatory factor for *GPAT4* activity^67,68^. Within hematological cancer cell lines, the coessentiality network is significantly restructured, with the *ACACA*/*FASN* module correlated with *SCD* in most backgrounds (r = 0.35, P < 10^−18^) but strongly anticorrelated in 36 blood cancer cell lines (r = -0.52, P < 10^−3^, **Figure 3e**). The magnitude of this change in correlation is ranked #8 out of 31 million gene pairs (see Methods). In contrast, *ACACA* and *FASN* are weakly correlated with *CHP1* in most tissues but strongly correlated in AML, with underlying covariation largely driven by the PS phenotype in FASTS cells (**Figure 3e**). This pathway sign reversal is confirmed in the single knockout fitness observed in our screens: *SCD* is strongly essential in MOLM13 but not in NOMO1 (**Figure 4e**).

Collectively these observations make a strong prediction about the metabolic processing of specific lipid species. Faster proliferation upon knockout of genes related to saturated fatty acid processing, coupled with increased dependency on fatty acid desaturase gene *SCD* (**Figure 5a**), suggests that these cells are at or near their carrying capacity for saturated fatty acids. To test this prediction, we exposed three FASTS cell lines and four other AML cell lines to various species of saturated and unsaturated fatty acids. FASTS cells showed significantly increased apoptosis in the presence of 200 µm palmitate (**Figure 5b-c**) while no other species of saturated or unsaturated fatty acid showed similar differential sensitivity. In addition, analysis of metabolic profiles of cells in the Cancer Cell Line Encyclopedia^69,70^ showed that saturated acyl chains are markedly overrepresented in triacylglycerol (TAG) in FASTS cells (**Figure 5d**), in contrast with other lipid species measured **(Supplementary Figure 7**). Palmitate-induced lipotoxicity has been studied in many contexts – and importantly, the role of *GPAT4* and *CHP1* in mediating lipotoxicity was well described recently^67,68^ – but, to our knowledge, this is the first instance of a genetic signature that predicts liposensitivity.

**Figure 5.**
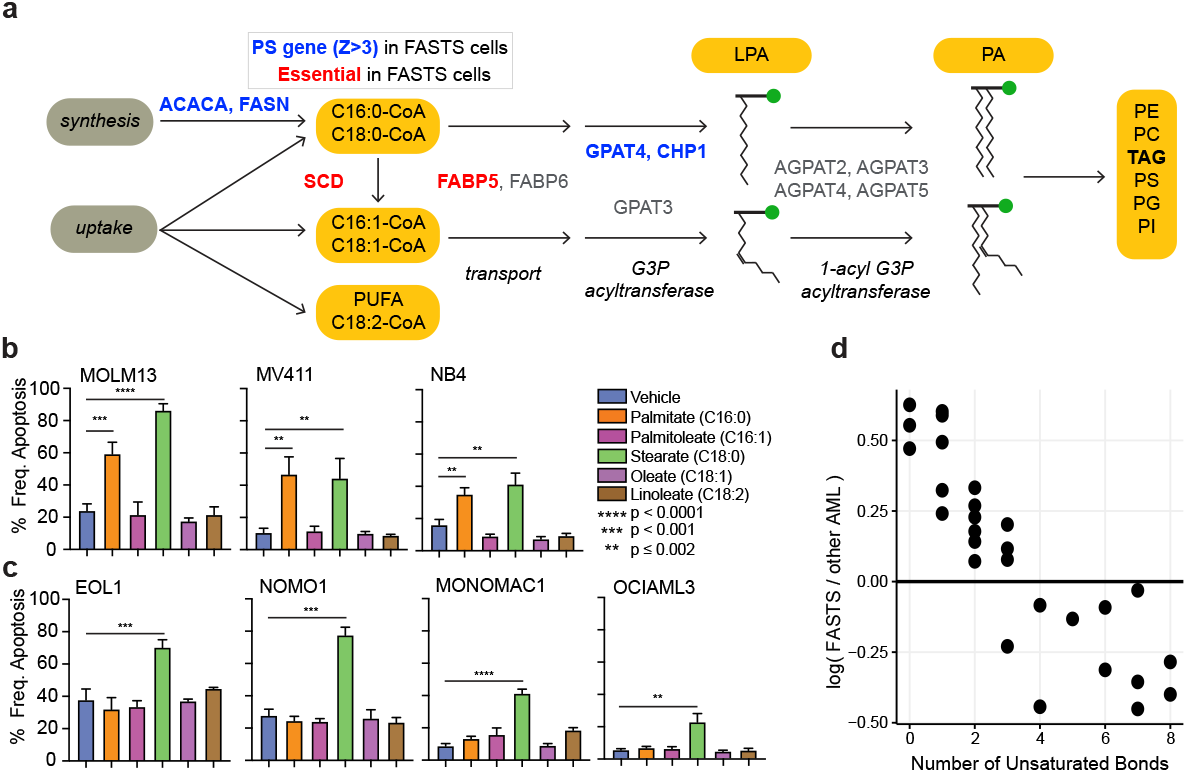
FASTS cells are sensitive to saturated FA. (a) Schematic of the fatty acid/glycerolipid synthesis pathway. Blue, PSG in FASTS cells. Red, essential genes. Pathway analysis suggests saturated fatty acids are a critical node. (b) Apoptosis of FASTS cells in response to media supplemented with 200 µm fatty acids. All three cell lines show marked sensitivity to palmitate. (c) Apoptosis of other AML cells in response to fatty acids shows no response to palmitate. (d) Triacylglycerol (TAG) species metabolite differences. The x axis represents the median difference of log10 normalized peak area of the metabolite in FASTS cells vs all other AML cells. The y axis represents the number of saturated bonds present. Each dot represents a unique metabolite.

### Prognostic signature for FASTS genes

To explore whether the FASTS phenotype has clinical relevance, we compared our results with patient survival information from public databases. Using genetic characterization data from CCLE^69^, we did not find any lesion which segregated FASTS cells from other CD33+ AML cells (**Figure 6a**), so no mutation is nominated to drive a FASTS phenotype *in vivo*. Instead, we explored whether variation in gene expression was associated with patient outcomes. We included genes in the core FASTS module as well as genes with strong genetic interactions with *ACACA/FASN* in our screen (**Figure 6a**). To select an appropriate cohort for genomic analysis, we first considered patient age. Although AML presents across every decade of life, patients from whom FASTS cell lines were derived are all under 30 years of age (sources of other AML cells ranged from 6 to 68 years; **Figure 6b**). With this in mind, we explored data from the TARGET-AML^71^ project, which focuses on childhood cancers (**Figure 6c**). Using TARGET data, we calculated hazard ratios using univariate Cox proportional-hazards modeling with continuous mRNA expression values for our genes of interest as independent variables. We observed that 4/7 FAS genes, *GPAT4, CHP1, PCGF1*, and *GPI*, show significant, negative hazard ratios (HR), consistent with a tumor suppressor signature (**Figure 6d**), and that no other gene from our set shows a negative HR. Indeed, when stratifying patients from the TARGET cohort with high expression of *GPAT4, CHP1, PCGF1*, and *GPI* (**Figure 6e**), we observe significantly improved survival (P-value = 0.001, **Figure 6f**). These findings are not replicated for *GPAT4, CHP1*, and *GPI* in the TCGA^72^ or OHSU^73^ tumor genomics data sets, possibly because they sample older cohorts (Polycomb group subunit *PCGF1* is observed to have a HR < 1 within the OHSU cohort, **Supplementary Figure 8a**). However, age is not generally associated with expression of genes in the FAS cluster in either cell lines or tumor samples (**Supplementary Figure 8**).

**Figure 6.**
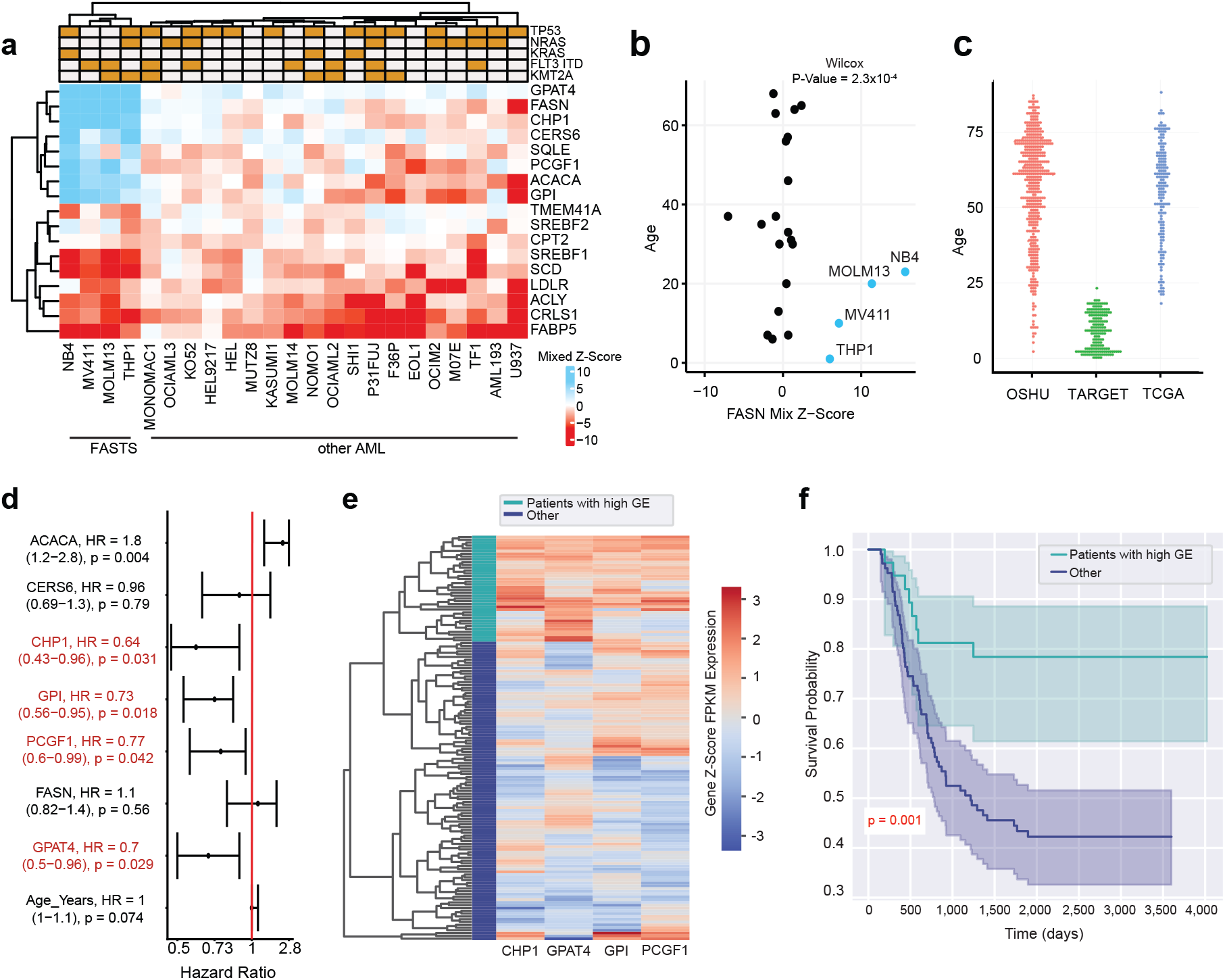
Prognostic signature of FAS module. (a) Heatmap of mixed Z scores for genes implicated in the genetic interaction network. Top, common AML lesions. (b) Mixed Z-score of FASN in AML cell lines vs. age of patient from which cell lines were derived. Blue, FASTS cells. (c) Age distribution of AML patients in three public tumor genomics cohorts. (d) Hazard ratios (95% CI; univariate Cox proportional hazards test) for expression of genes in (a), using genomics and survival data from TARGET. (e) Hierarchical clustering of gene expression in TARGET, using the four genes with negative HR. Green, high expression cluster. Blue, others. (F) Kaplan-Meier survival analysis of AML patients in TARGET, comparing patients in high expression cluster vs. others.

## Discussion

CRISPR screens have had a profound impact on cancer functional genomics. While research has been mainly focused on essential gene phenotypes, there is still much clinically relevant biology that can be uncovered by examining other phenotypes from a genetic screen. We establish a methodology that can reliably identify the proliferation suppressor phenotype from whole-genome CRISPR knockout genetic screens. This represents, to our knowledge, the first systematic study of this phenotype in the more than 1,000 published screens^8,10,11,13,48^.

The activity of proliferation suppressor genes is inherently context-dependent, rendering global classification difficult. As with context-dependent essential genes, the strongest signal is attained when comparing knockout phenotype with underlying mutation state. For example, wildtype and mutant alleles of classic tumor suppressor examples *TP53* and *PTEN* are present in large numbers of cell lines, enabling relatively easy discrimination of PS behavior in wildtype backgrounds, but most mutations are much more rare, reducing statistical power. Our model-based approach enables the discovery of PS phenotype as an outlier from null-phenotype knockouts. Using this approach, we recover COSMIC-annotated TSGs exhibiting the PS phenotype when wildtype alleles are expressed at nominal levels.

Co-occurrence of proliferation suppressors follows the principles of modular biology, with genes in the same pathway acting as proliferation suppressors in the same cell lines. We observe background-specific putative tumor suppressor activity for the PTEN pathway, P53 regulation, mTOR signaling, chromatin remodeling, and others. The co-occurrence network also reveals a novel module associated with glycerolipid biosynthesis, which exhibits the PS phenotype in a subset of AML cells. Analysis of the rewiring of the lipid metabolism coessentiality network in AML cells corroborated this discovery, and led us to define the Fatty Acid Synthesis/Tumor Suppressor (FASTS) phenotype in four AML cell lines. A survey of genetic interactions, using the enCas12a multiplex knockout platform, showed major network rewiring between FASTS and other AML cells and revealed strong genetic interactions in FASTS cells with *GPAT4*, a key enzyme in the processing of saturated fatty acids, and its regulator *CHP1*. Collectively these observations suggest that FASTS cells are near some critical threshold for saturated fatty acid carrying capacity, which we validated biochemically by treatment with fatty acids and bioinformatically through analysis of CCLE metabolomic profiles.

Confirming the clinical relevance of an *in vitro* phenotype can be difficult. No obvious mutation segregates FASTS cells from other AML cells, and with only four cell lines showing the FASTS phenotype, we lack the statistical power to discover associations in an unbiased way. However, by narrowing our search to strong hits from the differential network analyses, we found a significant survival advantage in a roughly age-matched cohort for *GPAT4* and *CHP1* overexpression. This finding points to a wholly novel tumor suppressor signature for our PSG module, though significant further study is necessary to determine whether this gene expression signature confers a similar *in vivo* metabolic rewiring and sensitivity to saturated lipids.

The combination of genetic, biochemical, and clinical support for the discovery of a novel tumor suppressor module has several implications. First, it provides a clinical signature that warrants further research as a prognostic marker as well as a potential therapeutic target. Second, it demonstrates the power of differential network analysis, and in particular differential genetic interaction networks, to dissect the rewiring of molecular pathways from modular phenotypes. And finally, it suggests that there still may be much to learn from data-driven analyses of large-scale screen data, beyond the low-hanging fruit of lesion/vulnerability associations.

## Acknowledgments

This research was performed in partial fulfillment of the requirements for the PhD degree from The University of Texas MD Anderson Cancer Center UTHealth Graduate School of Biomedical Sciences; The University of Texas MD Anderson Cancer Center, Houston, Texas 77030. WFL, MMc, MMo, and TH were supported by NIGMS grant R35GM130119. MC is supported by a Kopchick fellowship and Pauline Altman-Goldstein Foundation Discovery Fellowship. EK is supported by a grant from the Prostate Cancer Foundation. MD is supported by a Schissler Foundation fellowship. TH is a CPRIT Scholar in Cancer Research (RR160032), and is additionally supported by MD Anderson Cancer Center Support Grant P30 CA016672. WFL is supported by the American Legion Auxiliary Fellowship in Cancer Research. This work was supported by the Andrew Sabin Family Foundation Fellowship (TH). Flow cytometry was performed at MDACC’s Advanced Cytometry & Sorting Facility supported by the NCI Cancer Center Support Grant P30CA16672.

## Author Contributions

WFL performed all PS discovery analysis. MF, AG, AS performed genetic interaction screens and PD, MC performed bioinformatic analysis. WFL, MC, EK, and MD performed all other bioinformatic analysis. MMo and MMc performed lipid profiling experiments. JGD and TH supervised the research. WFL and TH drafted the manuscript and all authors edited it.

## Competing Interests

JGD consults for Agios, Maze Therapeutics, Microsoft Research, and Pfizer; JGD consults for and has equity in Tango Therapeutics. WFL is a former consultant for BioAge Labs, and has equity in Kronos Bio Inc.

## Supplementary Materials and Methods

### Code Availability

Mixed Z-scoring, analysis using scoring metric, co-occurrence network, and survival analysis was conducted in R version 4.0.4^74,75^. dPCC correlation analysis, including empirical calculations were conducted in Python 3.8.2^76^, using the packages SciPy^77^, NumPy^78^, Matplotlib^79^, and pandas^80^. Code is made available at: https://github.com/hart-lab/tsg_crispr_screen_survey/. R packages tidyverse^81^, data.table^82^, and knitr^83–85^ were used for figure generation, data manipulation, and general R functions; mixtools^86^, permute^87^, and PRROC^88,89^ were used for data simulations present in figures and evaluation; biomaRt^90,91^, and org.Hs.eg.db^92^ were used in integrating data types; cowplot^93^, ggbeeswarm^94^, annotate^95^, RColorBrewer^96^, ComplexHeatmap^97^, gplots^98^, ggpubr^99^, grid^75^, circlize^100^, ggthemes^101^, ggExtra^102^, patchwork^103^, and ggplot2^104^, were used for figure aesthetics and generation. R packages survival^105,106^ and survminer^107^ were used for survival analysis and figure generation. Analysis related to Kaplan Meier and patient stratification was done in python version 3.8.5^108^ using the packages pandas^80^, numpy^78^, and scipy^77^ for statistical functions and data manipulation, seaborn^109^, plotly^110^, and matplotlib^79^ for figure aesthetics and generation, and lifelines^111^ for both statistical analysis and figure generation.

Analysis of enCas12a multiplex genetic screens was conducted in R 4.0.0 and Python 3.8.3^112^. Code for this analysis is available at https://github.com/PeterDeWeirdt/FASTS. R packages tidyverse^81^ and tidygraph^113^ were used for data manipulation and ggraph^114^ was used for graph visualization. Python packages SciPy^77^, NumPy^78^, Matplotlib^79^, pandas^80^, statsmodels^115^, plotnine^116^ were used for analysis and visualization. The Custom package gnt^117^ was used to calculate genetic interaction scores and gpplot^118^ was used to generate point density plots.

### Processing DepMap Screen and CCLE Genomics Data

Raw read count data and a map of guide RNAs were downloaded from the DepMap database (www.depmap.org)^10,48^ and Project Score database (https://depmap.sanger.ac.uk/)^13^. Avana data version 2020q4^49^ was used for this analysis. To avoid genetic interaction effects, we discarded sgRNAs targeting multiple protein coding genes annotated as public or update pending in The Consensus Coding Sequence (CCDS, release 22)^119^. Gene names in the guide RNA maps of Avana and Project Score were updated using human gene information obtained from ncbi ftp. Then, read count data for each replicate was passed through CRISPRcleanR^120^ with location information of sgRNAs for the Avana CRISPR library based on GENCODE^121^ to correct depletion effects caused by copy-number amplification. Following this correction, each guide’s log_2_ fold-change was calculated. For Project Score data, we used only the gene location information of KY library v1.0 which is built in CRISPRcleanR. Normalized TPM RNA-seq data, copy number data, and mutation annotations for CCLE^69^ cells were also downloaded from DepMap. Ensembl gene id in RNA-seq data was converted to gene symbol using cross reference downloaded from Emsembl Biomart^122^.

### Mixed Z-Score Metric

Mixed z-score metric was generated using R version 4.0.4 base stat packages^75^ and the mixtools^86^ normalmixEM function. To calculate the mixed z-score, individual guide log_2_ fold-changes for each cell line were passed through the default settings of the normalmixEM function to fit two distinction normal distributions. Of the 808 cell lines passed through this analysis, 805 cell lines were able to converge with two distinction normal distribution following 1,000 iterations. The calculated mean and standard deviation of the higher (more positive) distribution were recorded. Along with the uncorrected original gene log_2_ fold-change, was used to calculate the corresponding mixed z-score. The original and mixed Z-score formula is as follows:

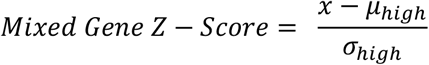

Where *x* is the original gene log_2_ fold-change, *μ_high_* is the average of the more positive fitted distribution, and *σ_hign_* is the standard deviation of the more positive fitted distribution. This metric was calculated for the DepMap 2020q4^49^ screen set, and the Sanger’s DepMap^13^ screen set for **Supplementary Figure 3**.

### Comparisons of Fitness Scoring Metrics

The following describes our comparative analysis of screening algorithms observed in **Supplementary Figure 1**. JACKS^43^ and BAGEL^41,42,123^, software was downloaded from their corresponding GitHub official distribution sites: https://github.com/felicityallen/JACKS, and https://github.com/hart-lab/bagel. We ran JACKS and BAGEL with raw fold change data of DepMap 2020q4 version^49^, gene guide map and replicate information. We obtained DepMap 2020q4 CERES scores from ‘dependency_score.csv’ downloaded from DepMap depository. Ranking was performed per screen and based on mean log_2_ fold-change values per gene.

We used the cancer gene census (CGC) list from COSMIC^45,46^ to compare fitness methods that can detect proliferation suppressor activity. Tumor suppressor genes (TSGs) from CGC represent a gene set of well-known proliferation suppressors. We separated the CGC gene list in two gene sets, genes with any tumor suppressor role in cancer representing true positive proliferation suppressor observations, and genes with any oncogene role in cancer representing false positives. Additionally, we added reference non-essential genes^7,47^ to the false positive list as these genes are not expected to demonstrate any phenotype. With these compiled lists, we evaluated each metric’s fitness scores, to see which metric would best separate the true and false positive gene lists. The R package PRROC was used for fitness scoring evaluation^88,89^.

### Direct Proliferation Suppressor Comparisons of Avana and Sanger Screen Datasets

The CRISPRcleanR^120^ corrected fold-change Sanger screen set^13^ was pushed through identical pipelines used to calculate the mixed z-score metric. Quality analysis of the mixed z-score metric for both data sets was pushed using identical gene sets described in the “Comparisons of Fitness Scoring Metrics” section. This analysis was restricted to only overlapping cell lines, 186 total, in both datasets.

The fitness enhancement introduced by PSG knockout, relatively weak compared to severe defects from essential gene knockout, often precludes detection in a shorter experiment. In the example F5 cell line (**Figure 1a**), a 2.5-fold change over a 21-day time course corresponds to a fitness increase of only ∼12% for rapidly growing cells, or a doubling time decrease from 24 to 21 hours. In a 14-day experiment, this increased proliferation rate would result in an observed log fold change of only ∼1.7, within the expected noise from genes with no knockout phenotype. This is explained in detail as follows:

#### Theoretical Fold-Change and Growth Rate Quantification

To assess hypothetical differences of proliferation suppressor fitness scoring metrics based on standard sampling times of screen collection taken from the Sanger and Avana databases^10,11,13,48^, we calculated theoretical cell population differences of wild-type and knocked out proliferation suppressor cell lines. The following formula can be used to calculate cell populations based on doubling rate per day:

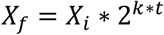

In this formula *X_f_* is the final population number of cells, *X_i_* is the initial population of cells, *k* is doubling time of the cells (in days), and *t* is time in days. In order to compare cells we can assume that these formulas are consistent with both wild-type cells and knocked out proliferation suppressor cells. With, knocked out proliferation suppressor cells the assumption is that these cells would grow faster compared to wild-type conditions and thus *k_ps_* > *k_wt_*, where *k_ps_* is the growth rate for proliferation suppressor knocked out cells, and *k_wt_* is the growth rate of wild type cells. These two independent growth rates are related as:

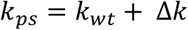

Δ*k* represents the change in growth rate resulting from genetic knockout, and is assumed to be positive. The growth rate formula for wild-type and proliferation suppressor cells is thus: 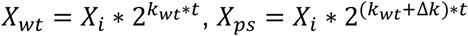

We then solved for Δ*k*, with *Log_2_(X_ps_/X_wt_)* as *Log_2_(FC)*, representing the fold-change difference between the cell populations at time *t*:

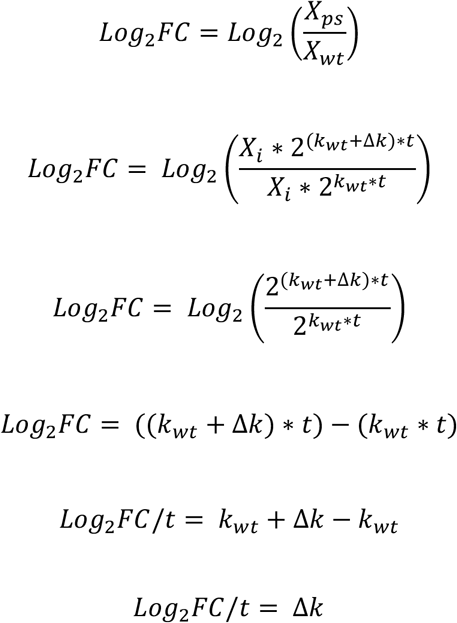

For a representative *Log_2_(FC)* of 2.5, which represents a sizable gain in fitness from a knocked-out proliferation suppressor, and *t* = 21 days, representing the time in which the Avana screens were sampled, we calculated Δ*k*:

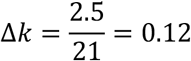

Using the calculated Δ*k* at 0.12, we can calculate the hypothetical *Log_2_(FC)* that would be expected at *t* = 14 days, representing the time in which the Sanger screens were sampled:

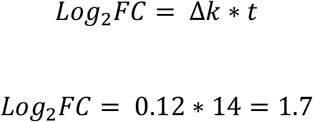

The resulting theoretical measurements demonstrate that Δ*k* can be identical between two samples, however the time in which the sample was taken will influence the ratio between the two measured cell populations. Taken together, this demonstrates that samples at shorter time points will demonstrate smaller quantified population size differences between wild-type and proliferation suppressor knocked out cells compared to samples taken at longer time points.

### Proliferation Suppressor Co-Occurrence Network

The co-occurrence network was developed based on FDR-corrected P-values from Fisher exact tests of all gene by gene comparisons that were identified as a proliferation suppressor more than once (584 genes total). Parallel processing, Fisher’s exact test, Benjamini & Hochberg FDR p-value adjustment were done using base R stat packages^75^. **Figure 2a** was created with heatmap.2 function from the R gplots^98^ package, with the dendrogram created through base R^75^ functions of euclidean distance, and complete agglomeration methods clustering of the Fisher’s exact test score between gene pairs. Smaller heatmaps displayed in **Figure 2c** were made using the R ComplexHeatmap library^97^. Network visualization was completed using Cytoscape^124^.

Network creation followed the corresponding steps; **1)** Identify all proliferation suppressor observations at a 10% FDR threshold (Z >= 3.83). **2)** Filter for gene proliferation suppressor observations that occurred at least 2 or more times, selecting for a total of 584 out of 18,111 genes available (3.2% total available genes); **3)** Create a binary (1 = proliferation suppressor, 0 = not proliferation suppressor) matrix of all 584 genes in all cell lines; **4)** Conducted Fisher’s exact test of every possible 2 x 2 contingency table of the 584 selected genes (n= 170,236 tests); and **5)** Adjust the corresponding p-values to FDR values, using a cutoff of 0.001 (0.1% FDR) to define edges. By assessing gene edges through Fisher exact-tests, we observe gene associations that are based on the relative proportion of co-occurrences between two genes.

### Proliferation Suppressor Network Enrichment

To test network enrichment of observed edges (**Supplementary Figure 4a**), we took 10,000 random samples of 462 (total number of edges in the co-occurrence network) gene pairs from the 170,236 available all by all gene pair Fisher’s exact test set. We then compared each sample to see the frequency of gene pairs observed to have some interaction within HumanNet^61^, excluding genetic interactions observed solely in the co-essentiality network component^21^ (generated from the same data) to prevent circularity. Additionally, we compared our selected mixed Z-Score cutoff against other various Z-Score cutoffs to ensure that we observed appropriate edge representation from HumanNet (**Supplementary Figure 4b**). Networks were made using identical pipelines and Fisher’s exact test set cutoffs with Z-Score cutoffs between 3 and 8 at 0.2 increments.

### Differential Pearson Correlation Coefficient Analysis

Differential Pearson correlation coefficient (dPCC) analysis was conducted to identify genetic fitness distinctions between AML cells and all other cells (**Figure 3**). Initial correlations (**Figure 3a**) of FAS cluster genes, PCGF1, CERS6, GPI, FASN, CHP1, GPAT4, and ACACA were calculated with R version 4.0.4 base stat packages^75^ and plotted in ggplot2^104^.

Following this observation, a follow up dPCC analysis was conducted on the FASTS cluster genes to assess dPCC quality. Cell line screens with low quality (Cohen’s D < 2.5 or recall of known core essential genes < 60%) were excluded, leaving 659 cell lines. Following this filtering step, two gene-by-gene correlation matrices were calculated. The first correlation matrix calculated all gene by gene pairs in only the available AML cell lines (n=17). The second matrix calculated all gene by gene pairs in the remaining 642 cell lines. The dPCC matrix is therefore the AML correlation matrix minus the non-AML correlation matrix.

Each gene-pair has a unique joint distribution of mixed Z scores; thus, the significance of each dPCC score must be calculated individually. To do this, we generated null distributions for dPCC for each gene pair. We took random selections without replacement of 17 cell lines (matching the n of AML cells), calculated all gene by gene pairwise correlations within this selection and within the remainder, and calculated dPCC. We repeated this sampling and calculation 1,000 times to generate a unique null distribution of dPCC for each gene pair and calculated an appropriate P-value for the observed dPCC above (right tailed for positive dPCC, left tailed for negative dPCC).

Genes which showed signficant knockout phenotype (|mixed Z| > 5) and AML-specific change in correlation (dPCC P<0.001) with a gene in the connected clique in the co-occurrence cluster (*CHP1, GPAT4, ACACA, FASN, GPI, CERS6, PCGF1*) were selected for further analysis (**Figure 3e**). **Figure 3e** was made using the R ComplexHeatmap library^97^. **Figure 3c-d** plots were made using the Python package Matplotlib^79^.

### Cell culture for Genetic Screens

MOLM13 and NOMO1 cells screened with the Cas12a-mediated genetic interaction library at the Broad Institute were obtained from the Cancer Cell Line Encyclopedia.

All cell lines were routinely tested for mycoplasma contamination and were maintained without antibiotics except during screens, when the media was supplemented with 1% penicillin/streptomycin. Cell lines were kept in a 37 °C humidity-controlled incubator with 5.0% carbon dioxide and were maintained in exponential phase growth by passaging every 2-3 days. The following media conditions and doses of polybrene, puromycin, and blasticidin, respectively, were used:

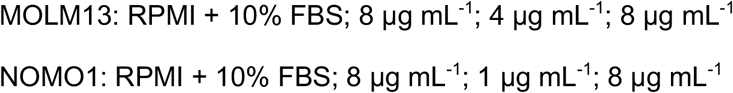

### Pooled screens

Cell lines stably expressing enCas12a (pRDA_174, Addgene 136476) were transduced with guides cloned into the pRDA_052 vector (Addgene 136474) in two cell culture replicates at a low MOI (∼0.5). Transductions were performed with enough cells to achieve a representation of at least 750 cells per guide construct per replicate, taking into account a 30–50% transduction efficiency. Throughout the screen, cells were split at a density to maintain a representation of at least 1000 cells per guide construct, and cell counts were taken at each passage to monitor growth. Puromycin selection was added 2 days post-transduction and was maintained for 5 days. 14 days and 21 days after transduction, cells were pelleted by centrifugation, resuspended in PBS, and frozen promptly for genomic DNA isolation.

### Genomic DNA isolation and PCR

Genomic DNA (gDNA) was isolated using the KingFisher Flex Purification System with the Mag-Bind® Blood & Tissue DNA HDQ Kit (Omega Bio-Tek #M6399-01) as per the manufacturer’s instructions. The gDNA concentrations were quantitated by Qubit. For PCR amplification, gDNA was divided into 100 μL reactions such that each well had at most 10 μg of gDNA. Per 96 well plate, a master mix consisted of 144 μL of 50x Titanium Taq DNA Polymerase (Takara), 960 μL of 10x Titanium Taq buffer, 768 μL of dNTP (stock at 2.5mM) provided with the enzyme, 48 μL of P5 stagger primer mix (stock at 100 μM concentration), 480 μL of DMSO, and 1.44 mL water. Each well consisted of 50 μL gDNA plus water, 40 μL PCR master mix, and 10 μL of a uniquely barcoded P7 primer (stock at 5 μM concentration).

PCR cycling conditions: an initial 1 min at 95 °C; followed by 30 s at 94 °C, 30 s at 53 °C, 30 s at 72 °C, for 28 cycles; and a final 10 min extension at 72 °C. PCR primers were synthesized at Integrated DNA Technologies (IDT). PCR products were purified with Agencourt AMPure XP SPRI beads according to manufacturer’s instructions (Beckman Coulter, A63880).

Samples were sequenced on a HiSeq2500 Rapid Run flowcell (Illumina) with a custom primer of sequence: 5’-CTTGTGGAAAGGACGAAACACCGGTAATTTCTACTCTTGTAGAT. The first nucleotide sequenced with the primer is the first nucleotide of the guide RNA, which will contain a mix of all four nucleotides, and thus staggered primers are not required to maintain diversity when using this approach. Reads were counted by alignment to a reference file of all possible guide RNAs present in the library. The read was then assigned to a condition (e.g. a well on the PCR plate) on the basis of the 8 nt index included in the P7 primer.

### Scoring Genetic Interactions

To score genetic interactions we used a custom python package, gnt^117^, available on the python package index. We use log-fold changes (LFCs) as inputs to the scoring pipeline. We define *Y_ij_* as the observed LFC of a guide pair *i*, *j* and 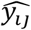 as this pair’s expected LFC. We then calculate the residual 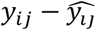 to generate an interaction score. To define expected LFCs, 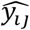 we fit a linear regression for each guide, *i*, saying

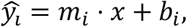

where *x* is the LFC of each guide individually and *m_i_* and *b_i_* are the fit slope and intercept for guide *i* (**Supplementary Figure 6b**). We refer to *i* as the anchor guide and its pairs as target guides. We then Z-score residuals within each anchor guide. This approach is similar to the one taken by Horlbeck *et al.*^33^.

To aggregate interaction scores at the gene level, we sum the z-scored residuals, *z_ij_*, for all constructs *i*, *j* targeting the gene pair *I*, *J*, fixing *I* as the anchor gene, and divide by the square root of the number of constructs targeting *I*, *J*. We repeat this calculation, fixing *J*as the anchor gene. We sum scores for both of these orientations and divide by 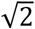 to arrive at a gene level Z-score.

### Cell Culture for Fatty Acid Response

Human cancer cell lines used at MD Anderson were obtained as follows: EOL1, MONOMAC1, NB4, OCIAML3 (DSMZ); MOLM13 and NOMO1 (Fisher); MV411 (ATCC). Identities were confirmed upon receipt and prior to experiments by STR typing (MDACC Characterized Cell Line Core). Absence of mycoplasma was confirmed monthly (Invivogen). All cell lines were grown at 37 °C in 5% CO_2_ in low attachment flasks (Greiner) and maintained at less than 1M cells ml^−1^. All but one line were cultured in RPMI-1640 with 25 mM HEPES (Sigma) supplemented with 10% FBS (Sigma), 2 mM Glutamax (Gibco), 1 mM sodium pyruvate (Gibco), 10,000 units ml^−1^ penicillin (Sigma), 10 mg ml^−1^ streptomycin (Sigma) and 100 µg ml^−1^ Normocin (Invitrogen). Complete medium was additionally supplemented with 0.1 mM non-essential amino acids (Gibco) for MONOMAC1.

### Fatty Acid Solutions

Fatty All chemicals were purchased from Sigma (St. Louis, MO). Solutions were prepared according to Luo *et al.*^125^ following best practices^126^. Fatty acid stock solutions were prepared in 100% ethanol at 50 mM for stearic acid or 200 mM for the rest. Fatty acid free bovine serum albumin (FAF-BSA) was dissolved in tissue culture grade (pyrogen free) water at 1.5 mM (10% w/v), filtered using 0.1 µm PES vacuum unit (Corning) and aliquoted for storage at -20°C. Ethanol stock solutions were diluted to 4 mM in FAF-BSA (molar ratio 2.7:1) and mixed gently at room temperature for 2 hours to facilitate conjugation. A vehicle control was prepared by diluting 100% ethanol in FAF-BSA to match the ethanol concentration in the 4 mM stearic acid solution. Vehicle or 4 mM solutions were aliquoted and stored at -80°C for up to 3 months. After thawing, aliquots were diluted 1:10 with complete medium to 400 µM, stored at 4°C and used within one week.

### Apoptosis Assay

Cells were seeded 24 hr prior to treatment in 500 µL complete medium in 24-well low attachment plates (Greiner) at 250,000 cells well^−1^. Quadruplicate wells received 500 µL FA working solution (400 µM) or vehicle (BSA+EtOH). Cells were treated at 200 µM for 48 hr. Treated cells were transferred to a deep 96 well plate and medium was discarded after centrifugation at 500 x g for 5 min. Cells were washed once with 1000 µL D-PBS (Sigma). Next, cells were resuspended in 300 µL binding buffer containing annexin-FITC and propidium iodide according to the manufacturer’s protocol (BD Biosciences) and transferred to a shallow 96 well V-bottom plate (Corning). After staining for 15 min at room temperature in the dark, cells were washed once with 300 µL binding buffer and finally resuspended in 100 µL binding buffer. Unstained and single stain controls were prepared for every cell line in a separate plate. Gates were adjusted such that 99% of unstained singlets fell below each threshold. See **Supplementary Figure 9** for complete gating strategy. Flow cytometry data were collected using a FACSCelesta analyzer equipped with an autosampler (BD Biosciences) and analyzed using FlowJo 10.5.3. Results shown are representative of three independent experiments conducted with sequential passages of each cell line.

### Metabolomics Analysis

This section describes the methods used within **Figure 5d** and **Supplementary Figure 7**. Metabolomics data acquired from Supplementary table 1 of Li *et al.*^70^ For analysis, normalized data (“1-clean data”) and coefficient of variation for each metabolite (“1-CV”) was used. Normalized data was filtered to select only AML cells that were present in Avana 2020q4^49^ screen set. Following filtering, the median of species present were taken, grouped by whether the measurement was from a FASTS AML or other AML cell line. The difference in median, representing the log ratio, was taken for each metabolite. Metabolites that had differences in medians less than the coefficient of variation were omitted from the plots. Acyl group and number of unsaturated bonds were obtained directly from the provided nomenclature.

### AML Patient Survival Analysis

This section describes the methods used within **Figure 6** and **Supplementary Figure 8 & 10**. The results published here are in part based upon data generated by the Therapeutically Applicable Research to Generate Effective Treatments (TARGET) initiative, phs000218, managed by the NCI. The data used for this analysis are available at dbGaP Study Accession: phs000465.v19.p8. Information about TARGET can be found at http://ocg.cancer.gov/programs/target.

Genes chosen for analysis were all genes shown to have an interaction with ACACA in **Figure 4h** and FASN. Gene annotations noted in the **Figure 6a** heatmap include any non-silent mutation, copy number loss for TP53 & KMT2A, and copy number gain for KRAS, NRAS, and FLT3. FLT3-ITD annotations were included in the FLT3 annotation row bar. Mutation annotations come from CCLE^69^, copy number calls come from the cBioPortal^127,128^ database, and FLT-ITD annotations come from the DSMZ catalogue^129^.

TARGET-AML^71^ data including age, genetic expression (HTseq FPKM UQ), time to event, and survival event outcomes, and TCGA^72^ patient ages and genetic expression were downloaded directly from the Xena^130^ database. The OHSU BeatAML^73^ age data was directly downloaded from the Vizome database, and genetic expression data was taken from the original publication. Age of patient derived cell lines were obtained from the Cellosaurus database^131^. Hazard ratios calculated from Cox proportional hazards modeling were done using the R survival^105,106^ package. Patient clustering stratification was done with clustering functions from the scipy package^77^, using Euclidean clustering and complete linkage settings. This output heatmap (**Figure 6e**) was created using functions from the seaborn^109^ package. We identified the patient cluster containing the highest overall expression of CHP1, GPAT4, GPI, PCGF1 from the heatmap using the fcluster function from scipy^77^. **Figure 6f** demonstrates the resulting survival comparison of the two patient clusters and was created with functions from the lifelines^111^ package, specifically, KaplanMeierFitter function for the Kaplan Meier curve, and the p-value reflecting the calculated logrank test of the two curves.

P-values related to schoenfeld tests calculated internally by the survminer package. For TARGET data analysis, patient expression profiles were chosen from primary tumor samples, filtering out samples from recurrent patients (42 such cases). Patient stratification is conducted based on stratifying patient groups into lower genetic expression (patients with genetic expression below the 75th percentile, n = 108), and higher genetic expression (patients with 75th percentile and above, n = 37). Computed hazard ratios for all tested genes within the TARGET cohort all passed the cox proportion hazards assumption (**Supplementary Figure 10**) by failing to reject the schoenfeld test null hypothesis.

## Supplementary Tables

**Table S1. Mixed Distribution Model Z-Score Matrix.** 808 cell line vs 18,111 gene matrix of mixed Z-score derived from log fold-change fitness scores.

**Table S2. COSMIC TSG PS Statistics.** Statistics of 116 COSMIC TSG genes when observed as a PS, vs other available data points. Includes number of times TSG is observed as a PS gene (count), mean and median TPM expression when observed as a PS gene and additional backgrounds (PS_Mean_Exp, Other_Mean_Exp, PS_Median_Exp, Other_Median_Exp), and non-silent mutation rate as a PS gene and additional backgrounds (PS_mut, Other_mut). Additionally includes a column of fisher’s exact test comparing mutated vs non mutated observations, and a Wilcox test comparing expression levels for each gene.

**Table S3. PSG Co-PS network.** Network of PSG co-occurrence observations related to **Figures 2c** and **S4c**, including fisher test metrics (p-value and FDR).

**Table S4. enCas12a Screen Gene Selection and Rationale.**

**Table S5. enCas12a Library Design.**

**Table S6. enCas12a Single Gene Knock-Out Measurements.** Z-score of mean Log fold-change.

**Table S7. enCas12a Double Gene Knock-Out Measurements.** Calculated Log fold-change and corresponding GI Scores for each gene pair.

## Supplementary figure legends

**Figure S1.**
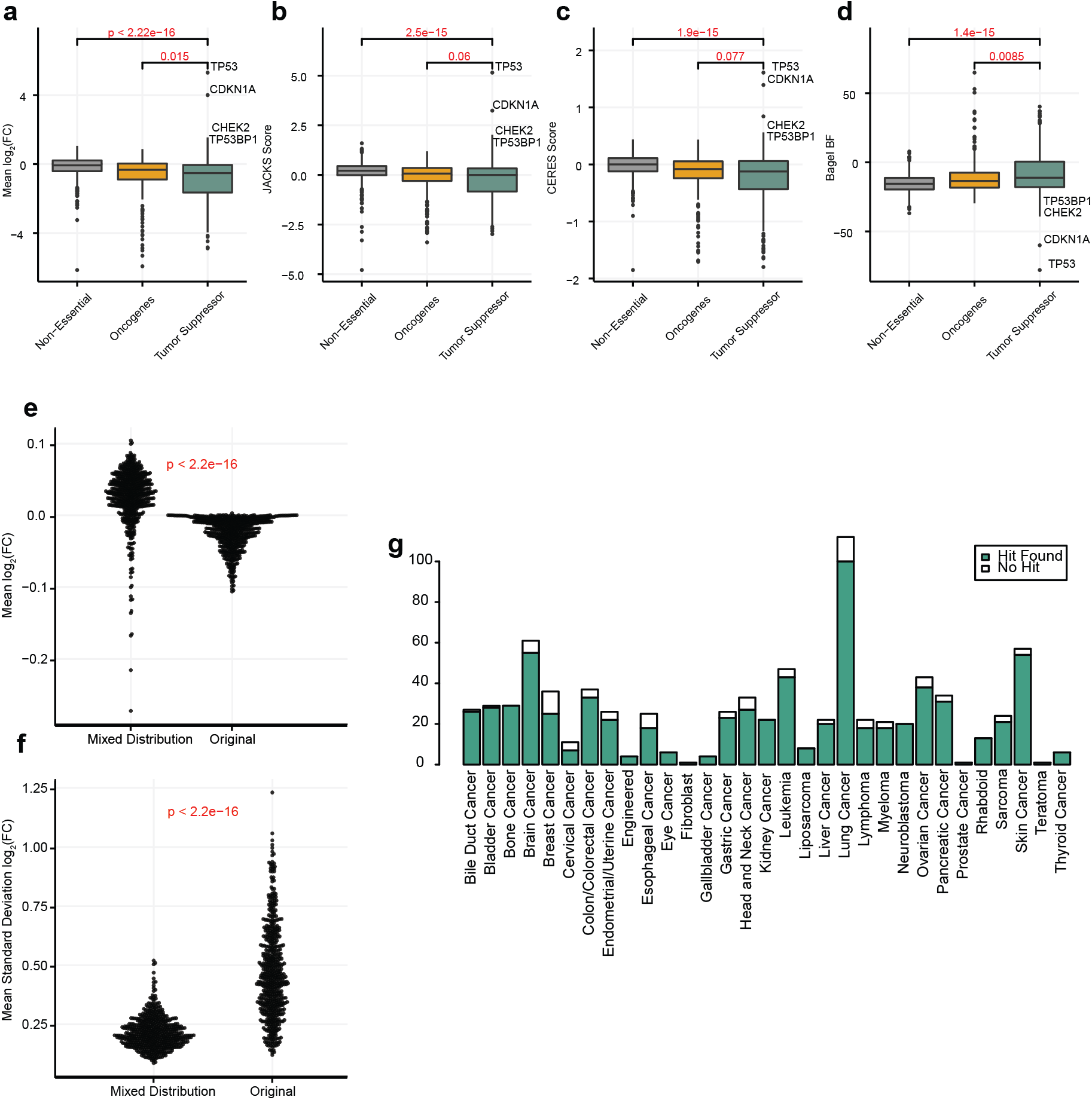
Discovery of Proliferation Suppressor genes extended. Fitness scoring distributions of non-essential genes, and non-overlapping COSMIC defined oncogenes and tumor suppressor genes; (a) mean log fold-change, (b) JACKS, (c) CERES, and (d) BAGEL. Selected screen for a-d matches the screen observed in Figure 1a. (e) Distribution of mean log fold-change of original distribution and mixed distribution. (f) Same (e) with mean standard deviation. (g) Bar chart by cell line lineage, where at least 1 PS gene at 10% FDR cutoff identified.

**Figure S2.**
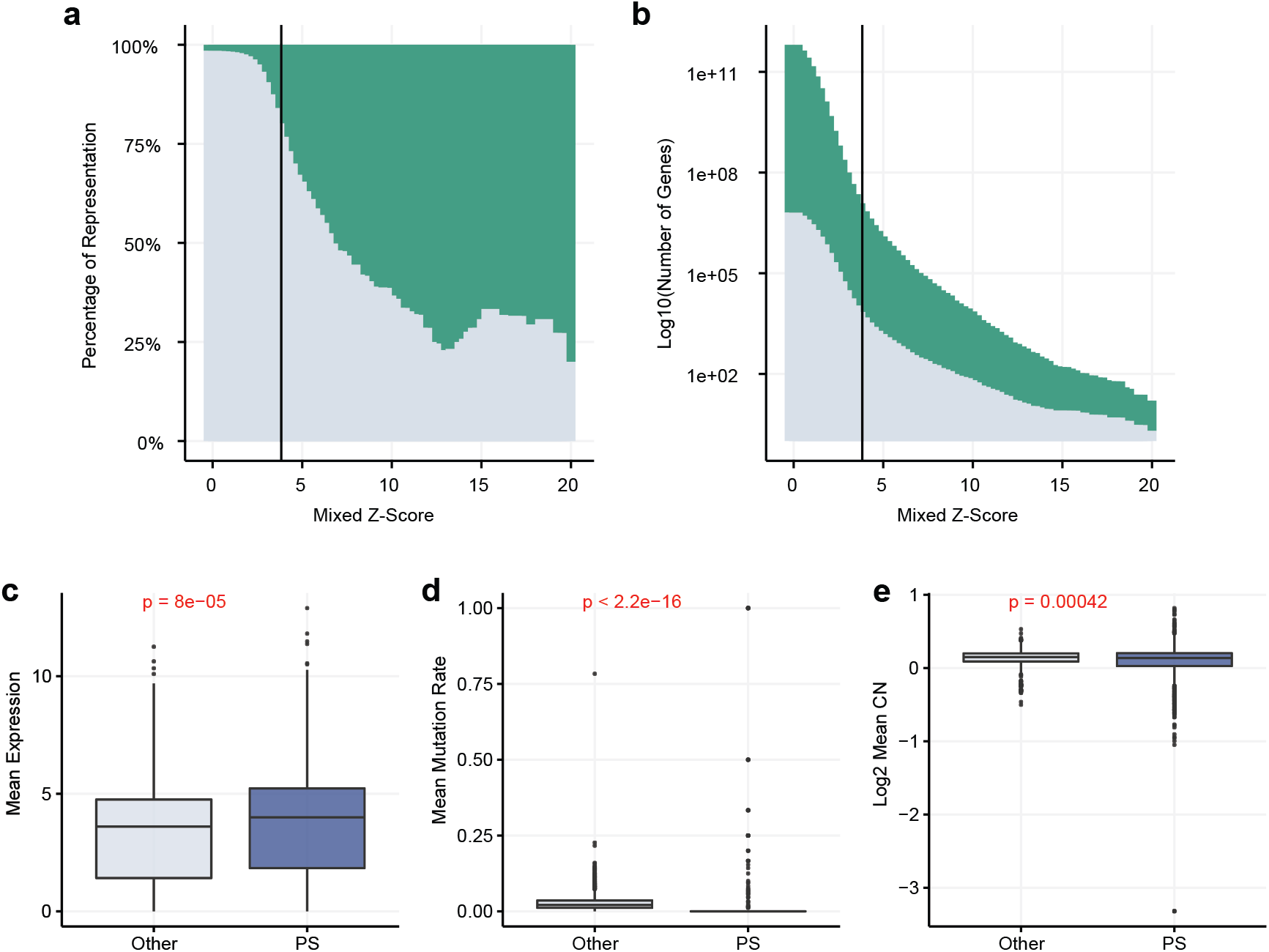
Proliferation Suppressor Gene Evidence. (a) Percent representation of COSMIC TSG (green) by corresponding label-shuffled Z-score. (b) Same as (a) with log10 y-axis of number of genes. (c) Mean TPM expression of PSG, grouped by PS observations (blue) vs every other available observation (gray) in which PSG were not observed as a PS. P value represents the corresponding Wilcoxon test. (d) same as (c) with mutation rate and (e) copy number.

**Figure S3.**
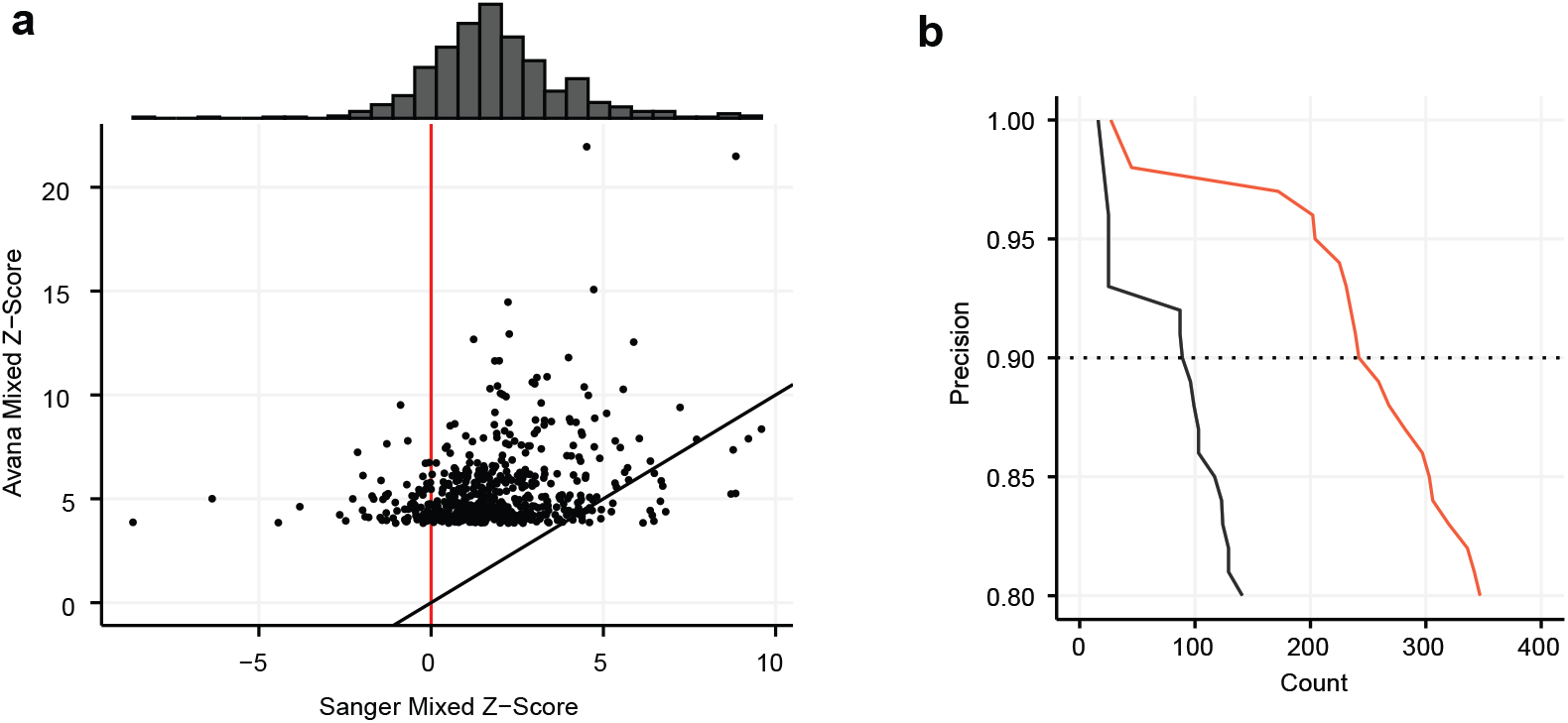
Avana vs Sanger Genetic Screens Comparison. (a) Precision vs. recall of mixed Z-score in matching screens from Avana (red), and Sanger (black). Dashed line, 90% precision (10% FDR). (b) Avana vs Sanger mixed Z-scores of genes identified as hits in Avana.

**Figure S4.**
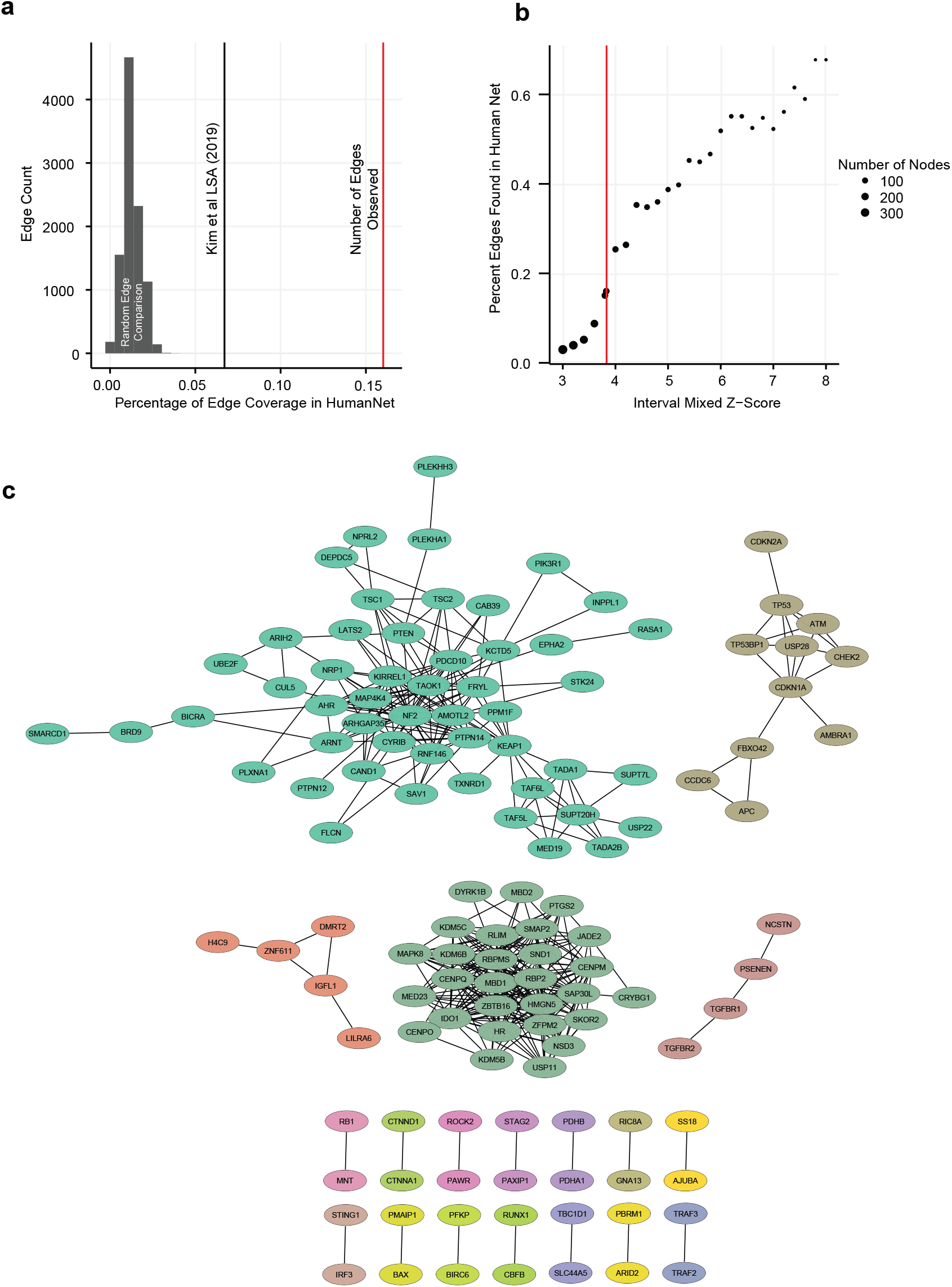
Co-occurrence of PS genes extended. (a) Empirical comparison of Co-PS network edges. Distribution represents random edges between genes identified in the network, and the percentage of edges identified in HumanNet with coessentiality network removed. Black line represents the percent of edges identified in the Kim *et al.* coessentiality network. Red line indicates the actual number of edges the Co-PS contains that are observed in HumanNet with coessentiality network removed. (b) Percent of edge coverage observed in HumanNet with coessentiality network removed against Co-PS edge FDR < 0.1%. networks at iterative label shuffled Z-score cutoffs. Red dot indicates actual cutoff used. (c) Remaining modules from the Co-PS network not included in Figure 2c.

**Figure S5.**
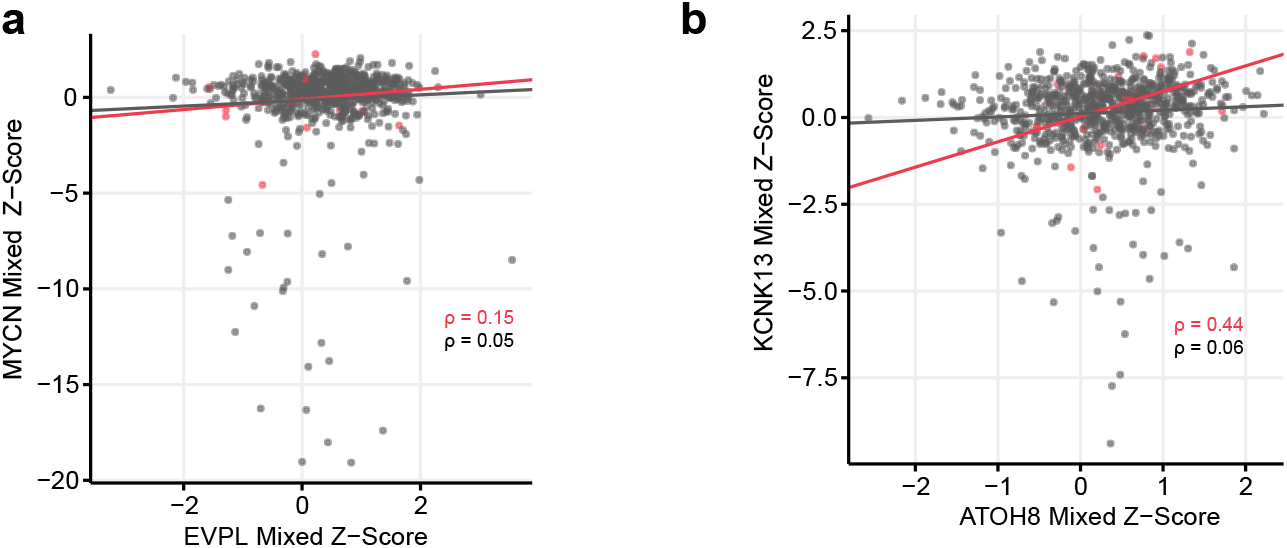
Examples of high dPCC resulting from data noise. (a) EVPL vs MYCN mixed Z-scores. Red indicates AML only observations, while gray indicates observations in all other cells. (b) same as (a) for ATOH8 vs. KNCK13 mixed Z-scores.

**Figure S6.**
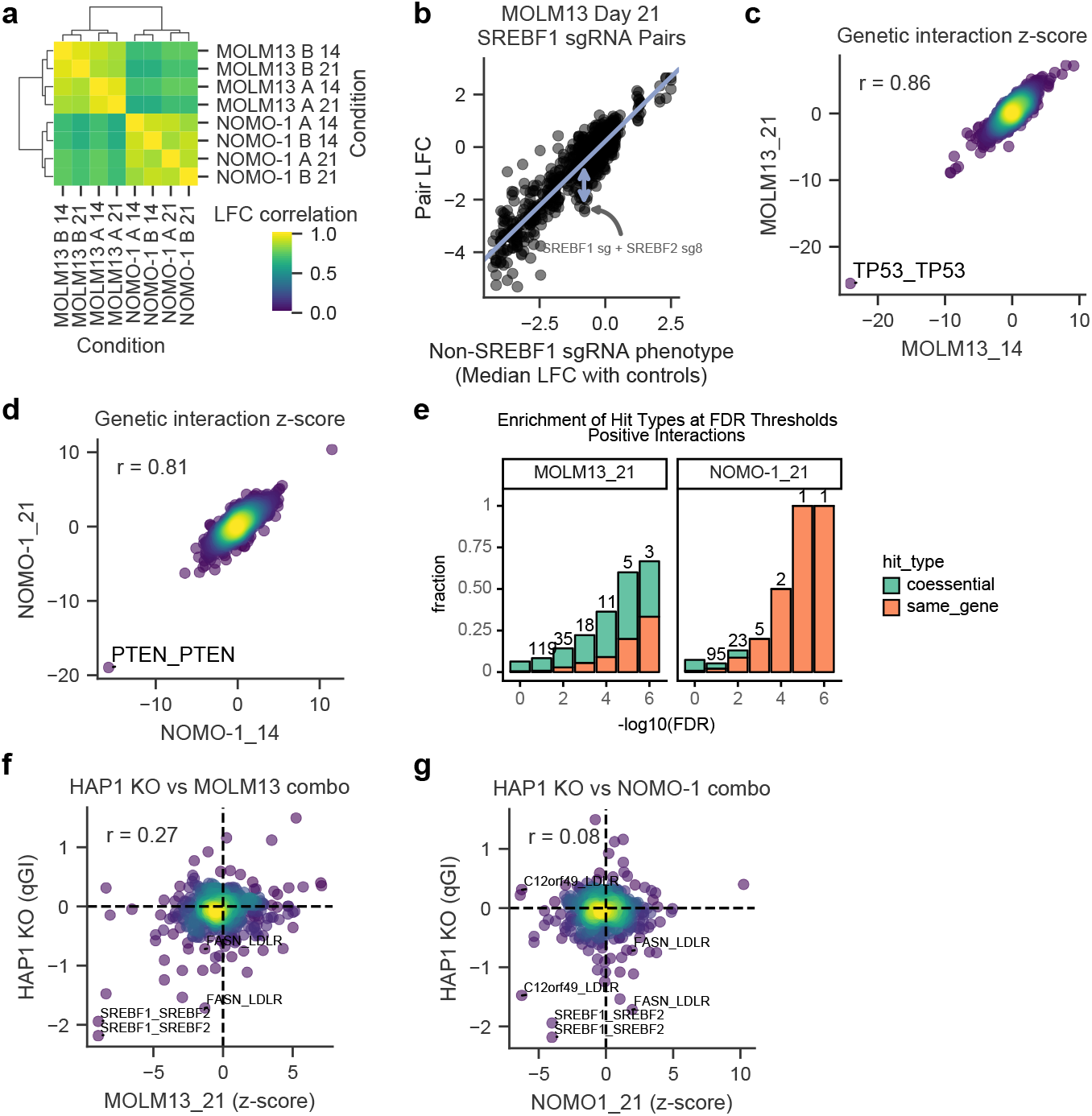
Combinatorial screen QC. (a) Replicate correlations. (b) Example calculation of residuals. (c) Correlation between genetic interaction scores for MOLM13. (d) same as (c) for NOMO1. (e) Fraction of coessential pairs or pairs that target the same gene at different FDR cutoffs for interactions with positive z-scores. (f) Comparison with qGI scores from Aregger *et al.* for MOLM13. (g) Same as (f) for NOMO1.

**Figure S7.**
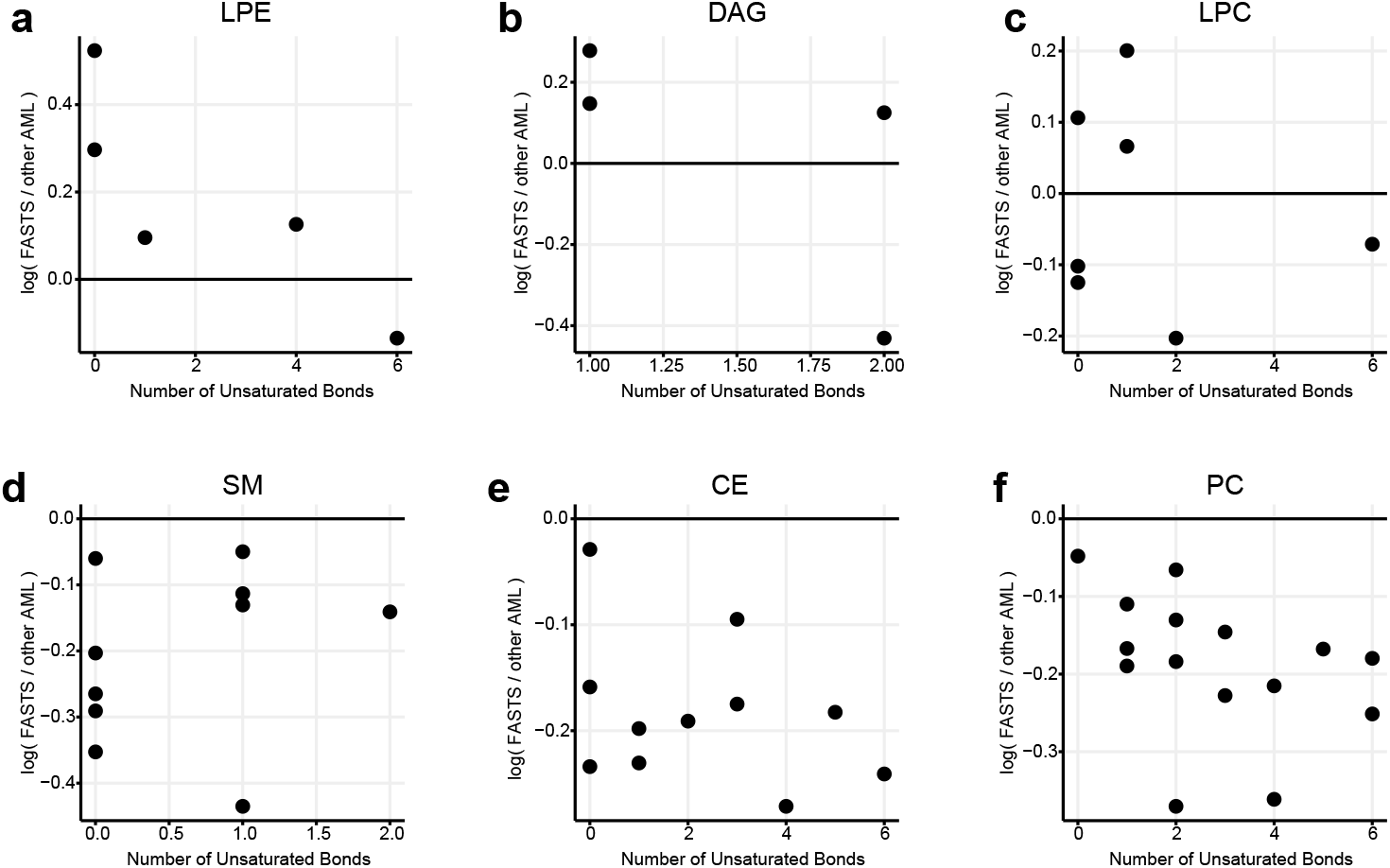
Additional metabolite comparisons. (a) Lysophosphatidylethanolamine (LPE) species metabolite difference. The x axis represents the median difference of log10 normalized peak area of the metabolite in FASTS cells vs all other AML cells. The y axis represents the number of saturated bonds present. Each dot represents a unique metabolite. (b) same for diacylglycerol (DAG), (c) lysophosphatidylcholine (LPC), (d) sphingomyelin (SM), (e) cholesterol ester (CE), and (f) phosphatidylcholine (PC) species.

**Figure S8.**
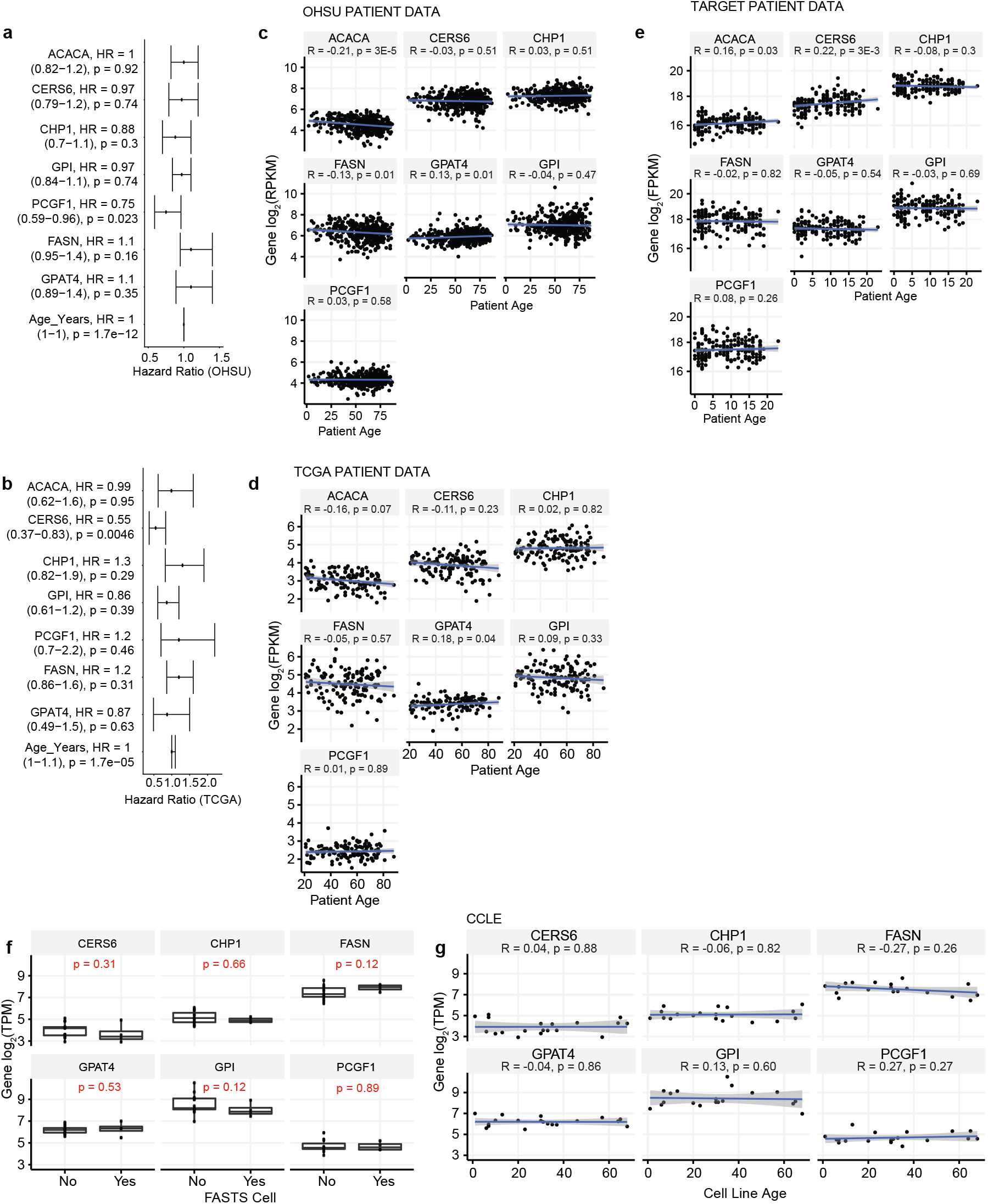
Comparisons of FAS genes against age in AML patient data. Hazard ratio calculations for FAS cluster genes in AML patient data coming from (a) OHSU - Tyner *et al.,* and (b) TCGA LAML. Spearman correlations of patient age against FAS gene expression in (c) OHSU, Tyner *et al.,* (d) TCGA LAML, and (e) GDC TARGET AML. (f) Boxplots of FAS gene expression in FASTS AML cell lines and non-FASTS AML cell lines from CCLE. (g) Spearman correlations of patient derived cell line age against FAS gene expression, coming from data in CCLE. ACACA is not included in (g) as it was not found in the CCLE expression data used in prior analysis.

**Figure S9.**
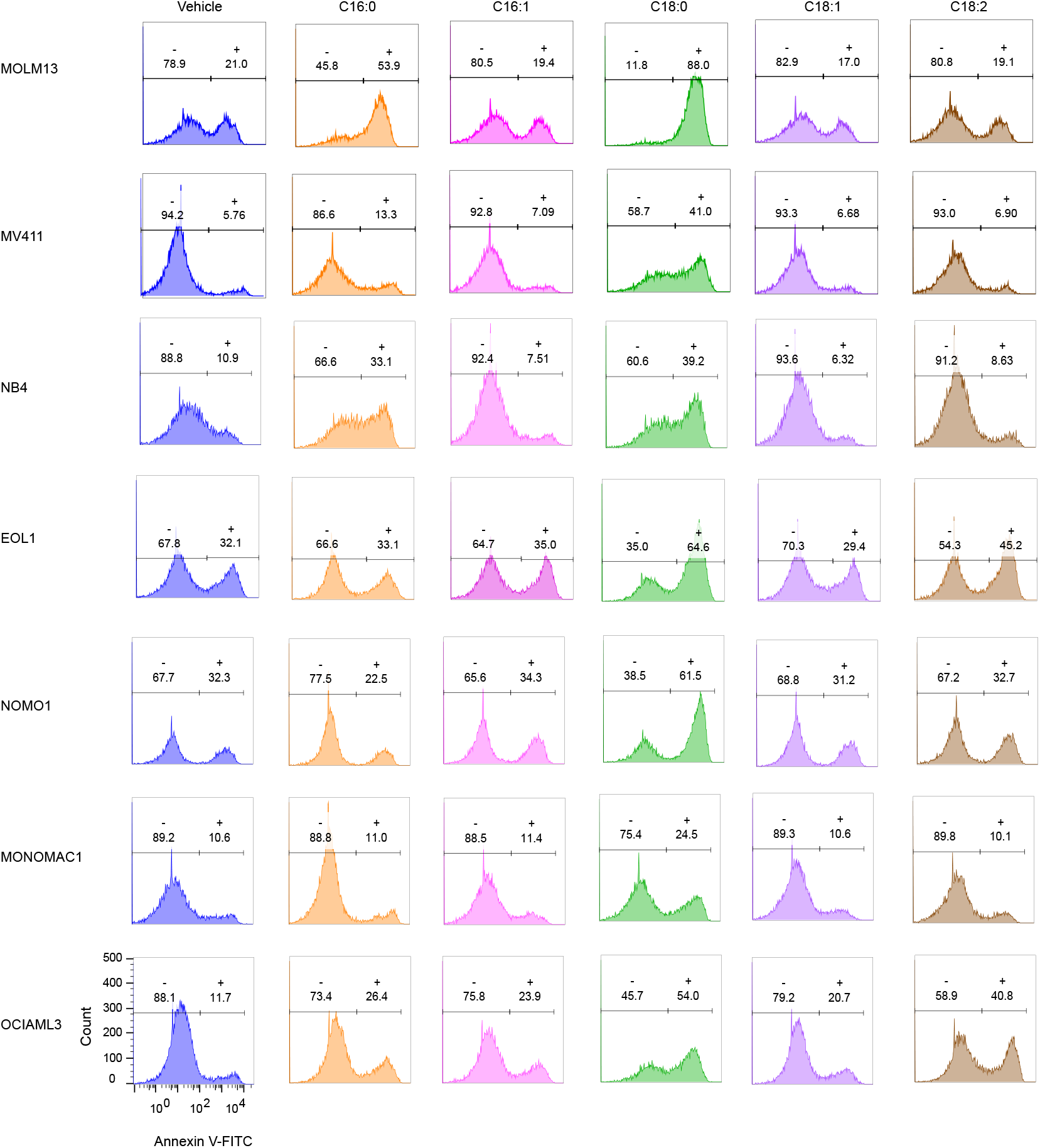
Sample flow cytometry plots. A representative flow cytometry data used to create bar graphs shown in figure 5b-c.

**Figure S10.**
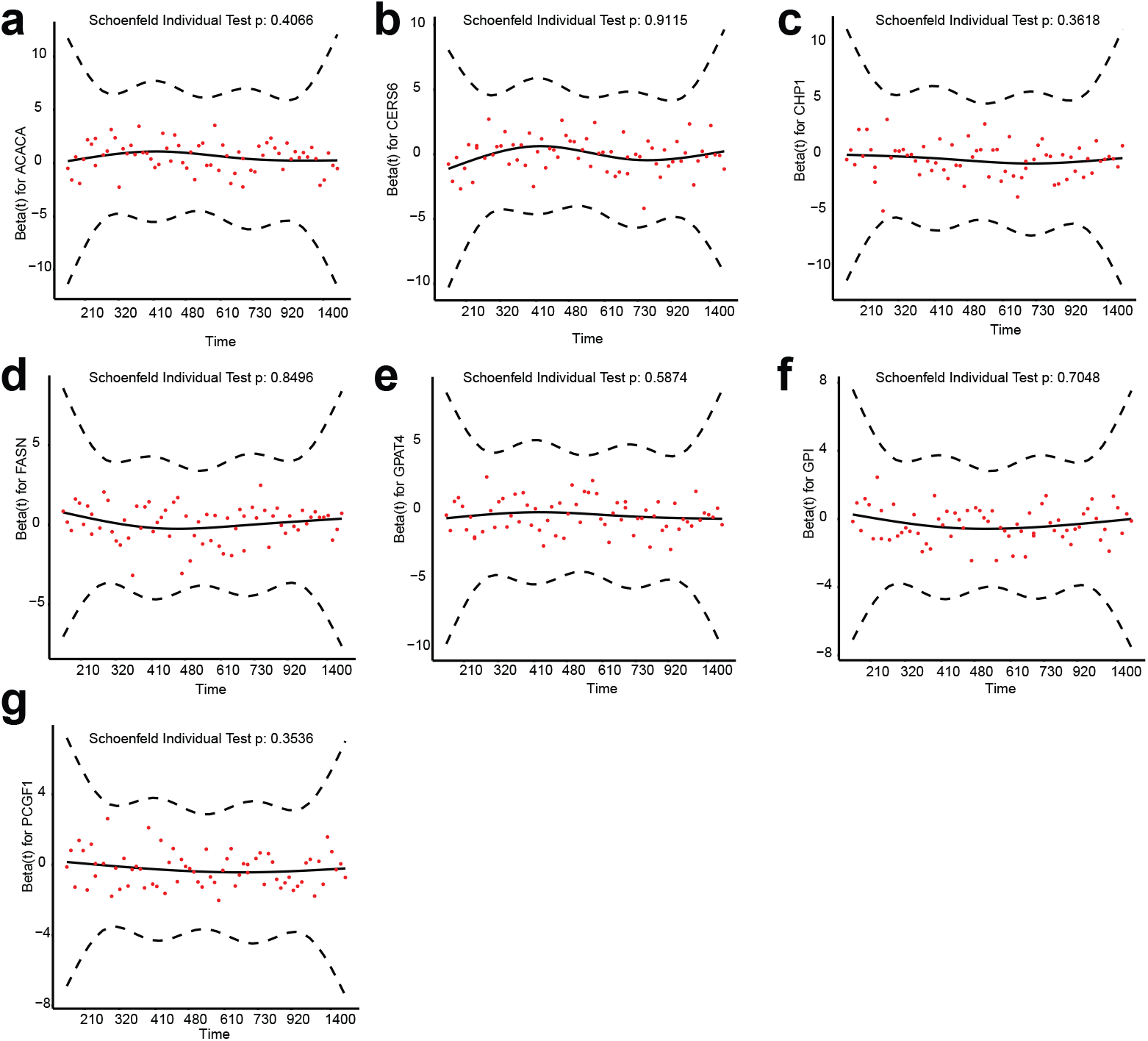
Testing the Cox Proportional Hazards Assumption. Assessing the Cox proportional hazards assumption with Schoenfeld tests of all genes in Figure 6d; (a) ACACA, (b) CERS6, (c) CHP1, (d)FASN, (e) GPAT4, (f) GPI, (g) PCGF1.

## Notes

### Competing Interest Statement

TH is a consultant for Repare Therapeutics. JGD consults for Agios, Maze Therapeutics, Microsoft Research, and Pfizer; JGD consults for and has equity in Tango Therapeutics. WFL is a consultant for BioAge Labs.

### Summary of Updates

In this revision, we update the algorithm for detecting proliferation suppressor genes to reflect Z-scores derived from a two-component Gaussian mixture model of the primary fold change data for each CRISPR screen, and the survival analysis is now based on clustering of patients with high expression of the putative tumor suppressor genes in the FASTS module.

